# Yeast Chd1p remodels nucleosomes with unique DNA unwrapping and translocation dynamics

**DOI:** 10.1101/376806

**Authors:** Jaewon Kirk, Ju Yeon Lee, Yejin Lee, Chanshin Kang, Soochul Shin, Eunhye Lee, Ji-Joon Song, Sungchul Hohng

## Abstract

Chromodomain-helicase-DNA-binding protein 1 (CHD1) remodels chromatin by translocating nucleosomes along DNA, but its mechanism remains poorly understood. Here, we employ a single-molecule fluorescence approach to characterize nucleosome remodeling by yeast CHD1 (Chd1p). We show that Chd1p translocates nucleosomes in steps of multiple base pairs per ATP. ATP binding to Chd1p induces a transient unwrapping of the exit-side DNA, and facilitates nucleosome translocation. ATP hydrolysis induces nucleosome translocation, which is followed by the rewrapping upon the release of the hydrolyzed nucleotide. Multiple Chd1ps binding to a single nucleosome sequentially moves a histone octamer with a preference to the center of DNA fragments, suggesting a new mechanism for regularly spaced nucleosome generation by Chd1p. Our results reveal the unique mechanism by which Chd1p remodels nucleosomes.

**Significance Statement:** There are four major ATP-dependent chromatin remodeler families: SWI/SNF, ISWI, CHD, and INO80/SWR1. The remodeling mechanisms of SWI/SNF and ISWI chromatin remodelers have been elucidated through extensive single-molecule studies, but it remains poorly understood how CHD chromatin remodeler operate. We use single-molecule FRET techniques, and show that Yeast CHD1 uses unique mechanisms to remodel a nucleosome.

## INTRODUCTION

Eukaryotic DNA is packaged hierarchically into a complex structure called chromatin. Chromatin is made up of fundamental units called nucleosomes, which are composed of 146 base pairs of DNA wrapped around a histone octamer (1, 2). DNA wrapped around histone cores to form nucleosomes is much less accessible to DNA-binding regulatory proteins (3). Cells, therefore, use ATP-dependent chromatin remodelers to modulate DNA accessibility during processes like DNA transcription, replication, and repair (4-7). There are four major ATP-dependent chromatin remodeler families: SWI/SNF, ISWI, CHD, and INO80/SWR1 (8). Most of these ATP-dependent chromatin remodelers form large multi-subunit complexes, each with a unique ATPase subunit that belongs to helicase superfamily 2 (5). In addition to this ATPase domain, each ATPase subunit contains several other distinct functional domains (9). For example, CHD family remodelers contain a chromo-domain that interacts with methyl-histone and/or DNA (5, 10-14). SWI/SNF family ATPases have C-terminal bromo domains that recognize acetylated histones (15, 16). The ATPase domain is thought to provide the mechanical force necessary for nucleosome remodeling, while the other domains in the ATPase subunit and other subunits in the complexes are thought to regulate the ways this mechanical force is applied in distinct situations and mechanisms. The mechanisms by which some of the chromatin remodelers—including ISWI, RSC, ACF, and CHD1—alter nucleosome structure have been explored with single-molecule technologies (17-33). In contrast to other chromatin remodelers for which the precise molecular mechanisms of nucleosome remodeling are relatively well studied, it remains still unclear how CHD1 remodels a nucleosome. Here, using single-molecule FRET (Fluorescence Resonance Energy Transfer) (34, 35), we unveil a novel mechanism by which yeast CHD1 (Chd1p) remodels nucleosomes.

## RESULTS

### Yeast Chd1p remodels nucleosomes with multiple base pair kinetic steps

ISWI and SWI/SNF family chromatin remodelers translocate nucleosomes in 1–2-bp steps (18, 19). A structural study on Chd1p also proposed 1-bp translocation mechanism for the CHD-family remodeler (36). However the actual step size of CHD1 has not been investigated. Therefore, we examined using single-molecule FRET (17) whether the same mechanism of 1-bp translocation is valid for Chd1p. We prepared nucleosomes using Widom 601 DNA sequence (37), labeled with Cy5 at the end of the DNA on the exit side, and Cy3 on H2A of the histone octamer. The Cy5 labeling site was selected to facilitate high levels of FRET before remodeling (Fig. S1A). We verified that fluorophore-labeling does not affect the remodeling activity of Chd1p (Fig. S1B). We then immobilized the nucleosomes on a polymer-coated quartz surface using the streptavidin-biotin interaction. We observed three peaks in the distribution of the FRET signal from the surface-immobilized nucleosomes, each corresponding to three different labeled species (Fig. S2). Of these three species, we used only the one with the highest levels of FRET in which Cy3 is attached to a position proximal to the exit side. This meant we were able to use FRET decrease to monitor the translocation of DNA at the exit side. After incubating the surface-immobilized nucleosomes with Chd1p, we added ATP to start nucleosome remodeling (Fig. 1A). Remodeling by Chd1p produced a stepwise reduction in the FRET signal (P1 and P2 in Fig. 1B), which suggests Chd1p remodels nucleosomes in a unidirectional manner. As previously observed for Chd1p as well as ACF (17, 21), we observed a recovery of FRET efficiency to its original value after an initial remodeling step (29% of remodeling events, Fig. S3A), which suggests that the nucleosome repositioning process can be reversed. At this moment, the nature of this revert of the remodeling is not clear. Although it is possible that this reversion might result from the Chd1p binding at the entry site, a consensus regarding how Chd1 binds to a nucleosome and remodels it has not been reached.

**Fig 1.**
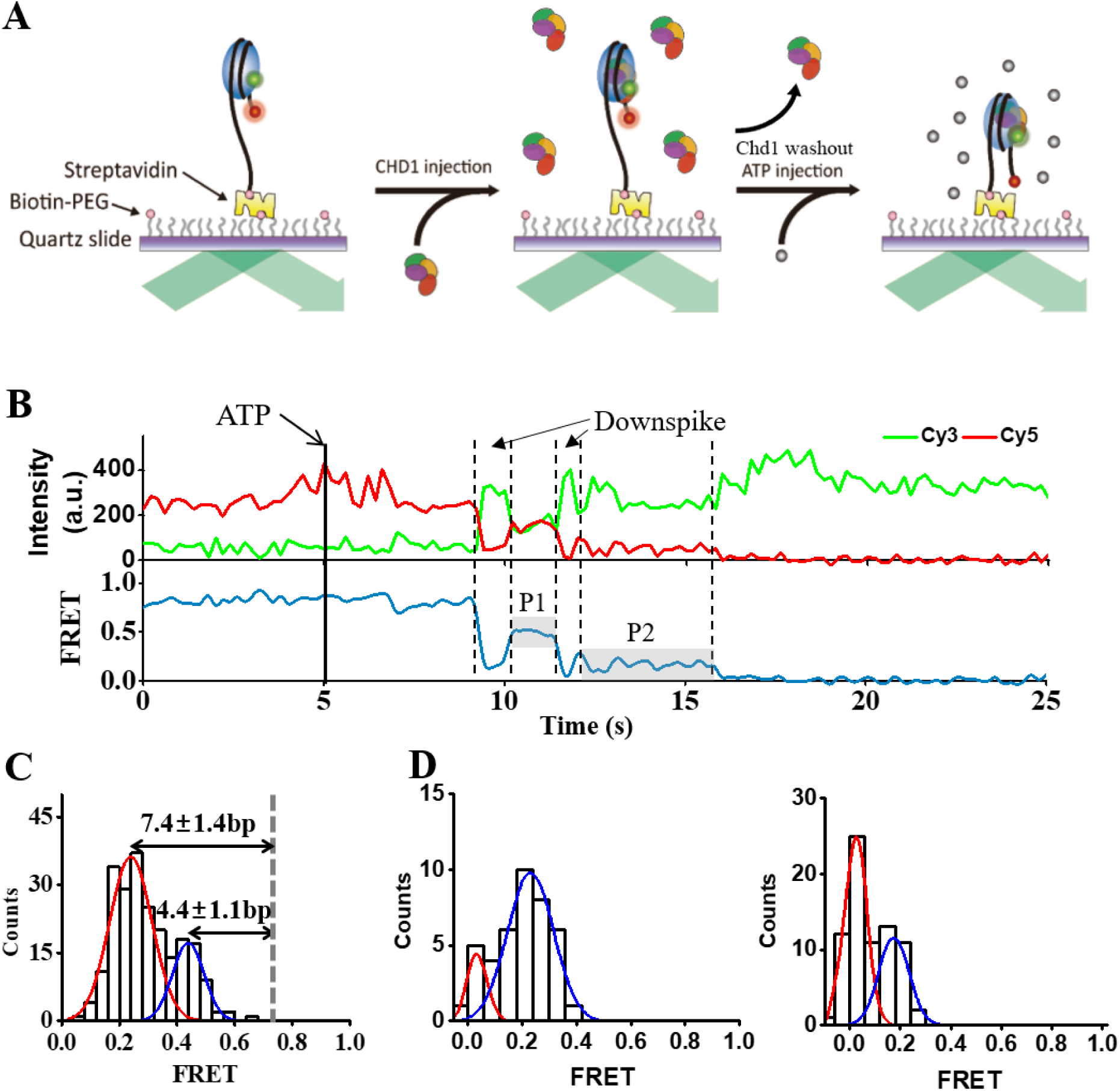
The kinetic step size of nucleosome remodeling by Chd1p. (A) Experimental scheme. Nucleosomes labeled with a FRET-pair (Cy3, green circle; Cy5, red circle) were immobilized on a microscope slide and incubated with Chd1p (180 nM). ATP was injected into the reaction chamber to initiate chromatin remodeling. (B) Representative fluorescence intensity (top, green for Cy3 and red for Cy5) and FRET (bottom) time traces during nucleosome remodeling by Chd1p. ATP was injected into the detection chamber at 5 seconds (solid line). This same color convention is used throughout the paper. (C) FRET histogram during the first translocation pause (P1). The histogram is fitted to two Gaussian functions. (D) FRET histograms during the second translocation pause (P2) after first translocations of a small step (left) or a large step (right). The histograms were fitted to two Gaussian functions. From 260 molecules with the initial high FRET state observed in five experiments, 224 molecules showed remodeling events (87.2 ± 5.3 %).

We were interested to observe two distinct populations in the FRET histograms after the first remodeling event (P1, Fig. 1C). Using FRET data calibrated with varying exit linker lengths (Fig. S4), we estimated that these two FRET populations correspond to small (4.4±1.1-bp) and large (7.4±1.4-bp) translocations. The second remodeling step (P2) also exhibited two FRET peaks, probably also corresponding to the small and large translocations (Fig. 1D). These occurred independently of the first remodeling step size (Fig. 1D), indicating that the step size is stochastically determined. These data imply that Chd1p may have a unique characteristic to other remodelers in its remodeling step size.

### Chd1p uses one ATP for each kinetic remodeling step

Other chromatin remodelers such as ISWI and RSC translocate nucleosomes with a 1 or 2-bp step per ATP molecule (18, 19). Although ISWI translocates nucleosomes with 3 or 7-bp kinetic steps, these kinetic steps comprise multiple 1-bp substeps (18). We, therefore, decided to examine whether the kinetic steps of Chd1p may also be composed of 1-bp substeps. First, we tried to observe the substeps by injecting the mixture of minimal ATP and saturating ATPγS (18). In contrast to ISWI chromatin remodelers which exhibited 1-bp substeps, we could not detect any substep (Fig. 2A), suggesting that the two—small and large—kinetic steps observed in Fig. 1 are fundamental remodeling step of Chd1p. In the same buffer condition, SNF chromatin remodeler clearly showed a 1bp substep (Fig. S5). However, this observation does not answer the question of how many ATPs are used for each fundamental remodeling step because multiple Chd1p may cooperatively function during the nucleosome remodeling. Chd1p is known to function as a monomer (38), but we found that multiple Chd1p can bind to a nucleosome in our experimental condition (Fig. S6). To observe the remodeling by monomeric Chd1p, we immobilized Chd1p using streptavidin-biotin interaction, and added a nucleosome labeled with a FRET pair (21). In this case Chd1p exhibited both the small (4.8±1.3-bp) and large (7.9±1.4-bp) steps as well (Fig. 2B), showing that the kinetic translocation steps of multiple base pairs are not due to multimerization of Chd1p. To confirm that single ATP is used for both the small and large remodeling steps, we preloaded nucleosome-Chd1p complexes with ATP in the absence of Mg^2+^. After washing free ATP from the reaction chamber, we added Mg^2+^ to start remodeling and then measured changes in FRET. Before the injection of Mg^2+^, free ATP was thoroughly washed out from the reaction chamber. Considering the efficiency of the buffer exchange system used (Fig. S7-9), it can be safely assumed that only prebound-ATP molecules could be used for nucleosome remodeling. Under this condition, the FRET histogram after the first remodeling step also showed two peaks corresponding to the small and large remodeling steps regardless of whether Chd1p or a nucleosome is immobilized (Fig. 2C-D). As expected from a monomeric Chd1p, when Chd1p was immobilized, only single remodeling steps were observed (Fig. 2C). Surprisingly, however, when a nucleosome was immobilized, we could observe multiple remodeling events (Fig. 2D), indicating that multiple Chd1ps binding to a single nucleosome can sequentially and unidirectionally remodel a nucleosome. In the case of multiple modeling events, the reversal of the nucleosome remodeling (Fig. S3B) was also observed with 33% probability.

**Fig 2.**
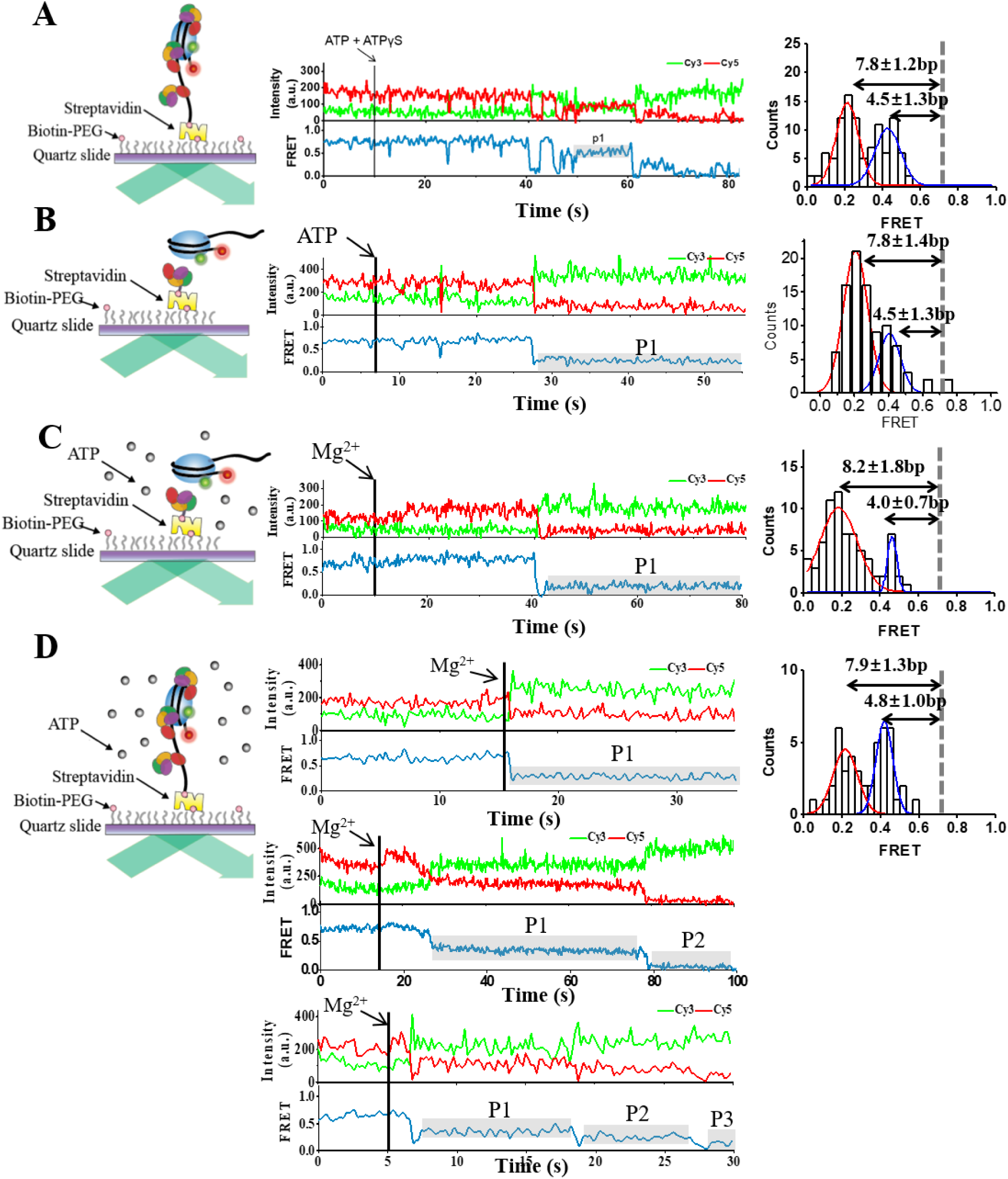
Nucleosome remodeling step size per ATP. Representative fluorescent intensity and FRET time traces (left), and FRET histograms during the first translocation pause (p1, right) for various experimental conditions: (A) Multiple Chd1ps are binding to a surface-immobilized nucleosome, and nucleosome remodeling is initiated by injecting 50µM ATP and 500µM ATPγS; (B) A nucleosome is binding to a surface-immobilized monomeric Chd1, and nucleosome remodeling is initiated by injecting 1mM ATP; (C) A nucleosome is binding to a surface-immobilized monomeric Chd1. The nucleosome-Chd1p complex is incubated with 100μM ATP and 10mM EDTA, and nucleosome remodeling is initiated by injecting 3mM Mg^2+^. (D) Multiple Chd1ps are binding to a surface-immobilized nucleosome. The nucleosome-Chd1p complex is incubated with 100μM ATP and 10mM EDTA, and nucleosome remodeling is initiated by injecting 3mM Mg^2+^. From the molecules that exhibited unidirectional translocation, percentages of one, two, and three remodeling events were 72.5%, 23.2%, and 4.3%, respectively. All FRET histograms are fitted to two Gaussian functions. The ratio of molecules that showed remodeling events to the total number of molecules with the initial high FRET state were 40.6 ± 2.1 % (104/256 in three experiments) for (A), 38.2 ± 7.6 % (92/240 in three experiments) for (B), 33.8 ± 2.3 % (73/215 in three experiments) for (C), and 12.0 ± 3.6 % (45/375 in eight experiments) for (D).

### ATP binding to Chd1p induces stochastic DNA unwrapping on the exit side

The FRET change during Chd1p-mediated nucleosome remodeling exhibited a characteristic, but not monotonic pattern; we consistently observed down-spikes in the FRET signal just before each translocation event (Fig. 1B). This suggests that the nucleosome structure is being altered right before nucleosome translocation. To better characterize the nature of these down-spikes, we performed similar remodeling experiments under different labeling schemes. First, we looked for Chd1p-induced conformational changes in the histone octamers using Cy3-H2A and Cy5-H4. In this case, we did not observe any appreciable change in the FRET signal (Fig. S10A), indicating that Chd1p does not induce large distance changes between the two labeling sites in the histone octamer. However, we cannot exclude a possibility that global conformational changes of the histone octamer in other positions. Next, to determine whether the down-spikes can be attributed to the dye labeling at specific positions on the histones, we moved Cy3 from H2A to H4 (Method). This had no effect on the down-spikes (Fig. S10B). The down-spikes coupled with a nucleosome remodeling were consistently observed when the dye labeling position on DNA was moved around the exit side (Fig. S10C-F). We, therefore, conclude that the FRET down-spikes are caused by substantial unwrapping of the DNA on the exit side prior to nucleosome translocation by Chd1p. Recent structural studies based on cryo-EM revealed DNA unwrapping in the presence of ADP-BeF3 (11, 20, 36). As DNA unwrapping occurs before DNA translocation, which requires ATP hydrolysis, we asked whether ATP binding itself is involved in the unwrapping that occurs on the exit side. We performed a double flow experiment by adding ATPγS followed by a step to wash the excess ATPγS from the reaction chamber (Fig. 3A). When we added ATPγS, we observed down-spikes without changes in the FRET signal corresponding to nucleosome translocation. This observation suggests DNA unwrapping occurs without ATP hydrolysis or nucleosome translocation. In some cases, we still observed the down-spikes for a while even after removal of ATPγS (Fig. 3A), suggesting that the binding lifetime of ATPγS to Chd1p can be tens of seconds, and the already bound ATPγS can induce unwrapping of exit side DNA several times. This retention of ATP analog by Chd1p explains why the Mg^2+^ flow assay described in figure 2C-D works. Consistently with the cryo-EM studies (11), we found that ADP-BeF3 induced stable DNA unwrapping (Fig. S11A). Interestingly, other ATP analogs, AMP-PNP and ADP-AlF4, did not induce any DNA unwrapping on the exit side (Fig S11B-C). To further investigate the relation between the binding of ATPγS and DNA unwrapping, we performed a kinetic analysis of the time delay between ATPγS injection and the first down-spike (t_i_), the down-spike dwell time (t_d_), and the high FRET dwell time after the first down-spike (t_h_) at varying ATPγS concentrations (Fig. 3B-D). The fact that all three of these kinetic parameters (t_i_, t_d_, and t_h_) can be nicely fitted to single exponential functions suggests each kinetic step has a single rate-limiting step. The ATPγS titration data showed that t_i_ and t_h_ depend on ATPγS concentration (Fig. 3E, Fig. S12A-B), whereas t_d_ seems to be independent of ATPγS concentration (Fig. 3F, Fig. S12C). Consistent with the observation that single ATPγS binding can induce multiple DNA unwrapping (Fig. 3A), t_h_ is shorter than t_i_ at low ATPγS concentrations. Thus, we conclude that DNA on the exit side is stochastically unwrapped by Chd1p while it is bound to ATP.

**Fig 3.**
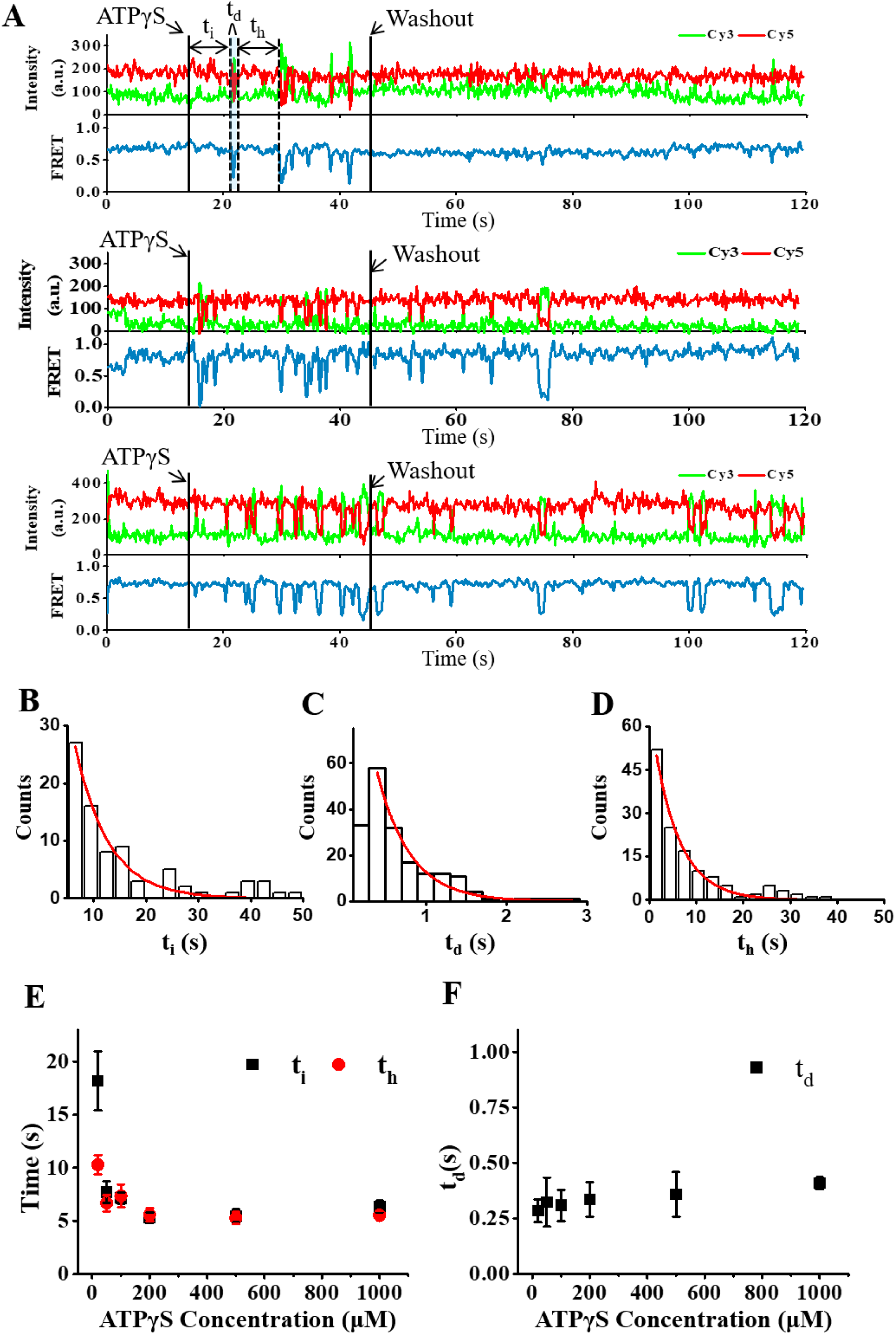
Stochastic unwrapping of the exit side DNA by Chd1p with ATPγS. (A) Representative fluorescence and FRET time traces showing FRET down-spikes upon addition of the ATP analog ATPγS (1 mM) at 30 seconds (solid line). In some molecules, the down-spikes remained for a while even after the removal of free ATPγS at 50 seconds (solid line). Kinetic parameters: t_i_, the delay between ATPγS injection and the first down-spike; t_h_, the dwell time between two adjacent down-spikes; t_d_, the dwell time of the down-spike. (B-D) Histograms of t_i_, t_h_, t_d_ at 1mM ATPγS, and their single-exponential fits with time constants of 6.35 ± 0.56 s for (B), 0.41 ± 0.02 s for (C), and 5.53 ± 0.41 s for (D) (red lines). (E) ATPγS titration of t_i_ and t_h_, the data were obtained by fitting the histograms to single-exponential functions (Figure S11A-B). (F) ATPγS titration of t_d_. The data were obtained by fitting the histograms to single-exponential functions (Figure S11C).

### Translocated DNA on the exit side is rewrapped after the dissociation of phosphate and ADP

To further characterize how ATP is used during Chd1p-mediated nucleosome remodeling, we analyzed, at varying ATP concentrations, the delay between ATP injection and the first down-spike (t_i_), the dwell time for down-spikes uncoupled to nucleosome translocation (t_du_), and the dwell time for down-spikes coupled to nucleosome translocation (t_dc_) as defined in Fig. 4A. We noticed FRET down-spikes frequently occurring just before remodeling, but remodeling did not necessarily always follow a down-spike. Therefore we distinguished down-spikes (dc) coupled to translocation from down-spikes (du) uncoupled to translocation. The t_i_ data are well-fitted to a single exponential function (Fig. 4B). Interestingly, the distributions of t_du_ and t_dc_ were clearly different from one another (Fig. 4C-D). The t_du_ data are reasonably well-fitted to a single-exponential distribution, while the t_dc_ data are well-fitted to a gamma distribution. The fact that the t_dc_ data cannot be fitted to a single exponential distribution suggests that multiple steps are involved in the kinetics of DNA rewrapping when it is coupled to nucleosome translocation. In addition, while t_i_ depended on ATP concentration (Fig. 4E, Fig. S13A), the t_du_ and t_dc_ data are independent of ATP concentration (Fig. 4F, Fig. S13B-C), supporting the conclusion that these events occur in the ATP-bound state. To determine whether the status of the hydrolyzed nucleotide after nucleosome translocation affects the duration of DNA unwrapping, we performed vanadate (39) and ADP titration experiments. At high vanadate or ADP concentrations, we observed increases in t_dc_, whereas t_du_ remained unaffected (Fig. 4G-H, Fig. S14). These data suggest that the DNA unwrapping state coupled to nucleosome translocation is maintained as long as the hydrolyzed nucleotide remains bound to Chd1p probably after nucleosome translocation. Then, upon phosphate and ADP dissociation, the translocated DNA is rewrapped. We confirmed that the pause between remodeling events (t_p1_ in Fig. 4A) depends on ATP concentration (Fig. S15), supporting the expectation that a new ATP hydrolysis cycle is required for each remodeling step.

**Fig 4.**
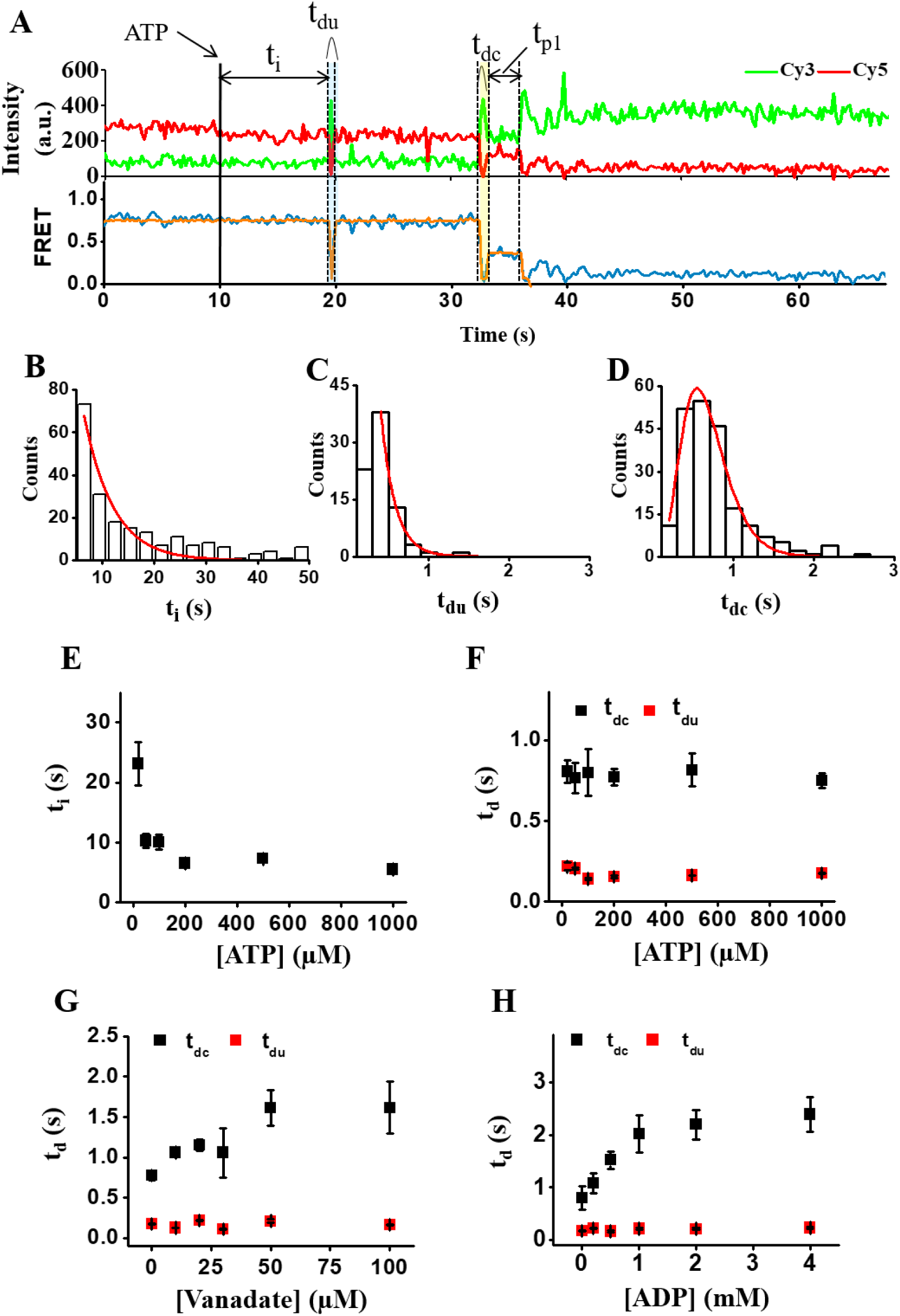
Rewrapping of the exit side DNA after phosphate and ADP dissociation. (A) Representative fluorescence intensity (top) and FRET (bottom) time traces reporting the translocation of the nucleosome after ATP injection at 10 seconds. Kinetic parameters: t_i_, the time delay between ATP injection and the first down-spike; t_du_, the dwell time of down-spikes uncoupled to translocation; t_dc_, the dwell time of down-spikes coupled to translocation. (B) A histogram of t_i_ at 1mM ATP and its fit to a single-exponential function with a time constant of 5.53 ± 0.49 s (red line). (C) A histogram of t_du_ at 1mM ATP and its fit to a single-exponential function with a time constant of 0.18 ± 0.04 s (red line). (D) A histogram of t_dc_ at 1mM ATP and its fit to a gamma distribution with n = 5.32 ± 0.17 and τ = 0.10 ± 0.02 s. (E) ATP titration of t_i_. The data were obtained by fitting the histograms to single-exponential functions (Fig. S13A). (F) ATP titration of t_dc_ (black) and t_du_ (red). t_du_ data were obtained by fitting the histograms to single-exponential functions (Figure S12B). t_dc_ is the mean ± SEM of those histograms (Fig. S13C). (G) Vanadate titration of t_dc_ (black) and t_du_ (red) at 1mM ATP. t_dc_ is the mean ± SEM of those histograms (Fig. S14C). t_du_ data were obtained by fitting histograms to single-exponential functions (Fig. S14D). (H) ADP titration of t_dc_ (black) and t_du_ (red) at 100µM ATP. t_dc_ is the mean ± SEM of those histograms (Fig. S14E). t_du_ data were obtained by fitting histograms to single-exponential functions (Fig. S14F).

### No time delay is observed between the exit and entry side remodelings

Cryo-EM studies revealed DNA unwrapping in the presence of ADP-BeF_3_ (11, 20, 36). Single-molecule FRET study showed that remodeling by ISWI proceeds with a time delay between the entry and exit sides (18). We therefore asked whether DNA unwrapping on the entry side and a time delay exist during Chd1p-mediated remodeling using nucleosomes labeled at the entry side (Fig. S1A). In contrast to what we observed with labeling at the exit side, we observed frequent FRET down-spikes even when Chd1p alone was injected without ATP (Fig. 5A). When we added ATP to initiate nucleosome remodeling, as expected from the results of the exit side remodeling (Fig. 1-2), we observed remodeling events corresponding to small and large remodeling steps (Fig. S16A-B). Interestingly, repetitive down-spikes were observed before ATP injection only in some remodeled molecules (top, Fig. 5B), but not in others (bottom, Fig. 5B). We do not currently understand the origin of the heterogeneity. When we did observe the down-spikes before remodeling, they became less frequent upon ATP injection (top, Fig. 5B; Fig. S16C). We also tested the degree to which the down-spikes were coupled to the translocation events and found that coupling on the entry side is significantly less than the exit side (22% vs. 77%, Fig. 5C). In case of the exit side remodeling, the coupling efficiency of the down spike and remodeling was apparently higher than the probability for a down-spike to occur simultaneously with nucleosome remodeling by chance, but the difference between them was not significant in the case of entry side remodeling (Fig. 5C). The experiment with a DNA substrate that has a 601 sequence in the reverse direction also gave the same result (Fig. 5D), indicating that the asymmetry of the 601 sequence is not the cause of the difference. Consistently with the cryo-EM studies (20, 36) and previous single-molecule FRET study (20), DNA on the entry side was unwrapped in the presence of various ATP analogs (Fig. S17).

**Fig 5.**
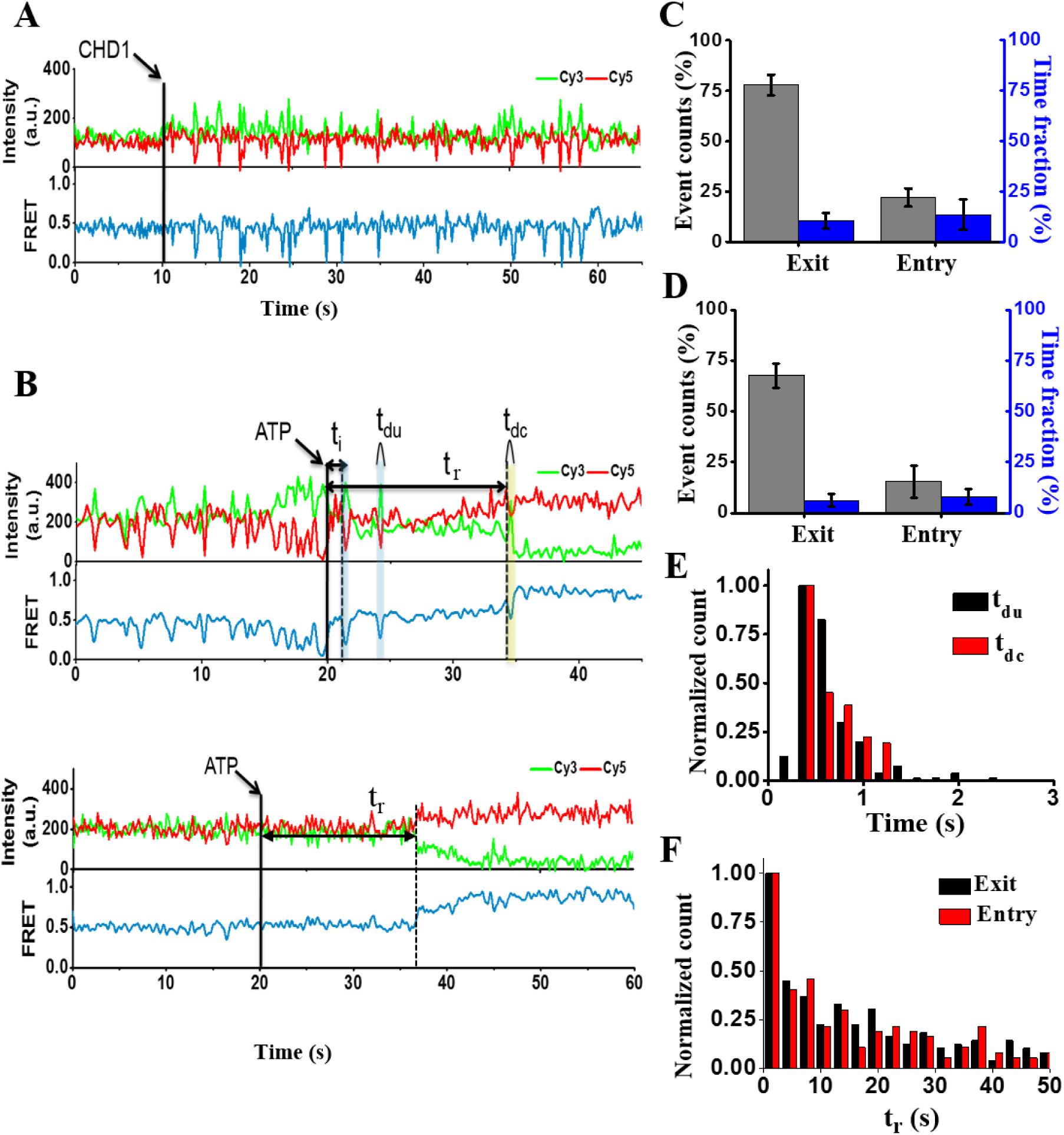
Entry-side remodeling by Chd1p. (A) Representative fluorescence intensity and FRET time traces for the entry-side labelled nucleosome in the presence of Chd1p. Chd1p was added at 10 seconds (solid line). (B) Representative fluorescence intensity (top) and their corresponding FRET (bottom) time traces showing remodeling on the entry side. Two different behaviors were observed: (upper panel) molecules showing down-spikes before ATP injection (85%) and (lower panel) molecules showing no down-spikes before ATP injection (15%). Kinetic parameters: t_i_, the delay between ATP injection and the first down-spike; t_du_, the dwell time of down-spikes uncoupled to nucleosome translocation; t_dc_, the dwell time of down-spikes coupled to nucleosome translocation; t_r_, the time between ATP injection and translocation. (C) The coupling efficiency of down-spikes with translocation on the exit (*N* = 237) and entry sides (*N* = 162). The data are shown as means ± SEM of 3 independent experiments. (D) The coupling efficiency of down-spikes with translocation on the exit (*N* = 120) and entry sides (*N* = 93) with the 601 sequence in the reverse direction. The data are shown as means ± SEM of 3 independent experiments. (E) Histograms of t_du_ (black) and t_dc_ (red) of the entry-side labelled nucleosome, in presence of 1mM ATP. (F) Histograms of t_r_ on the exit-side labelled nucleosome (*N* = 237) and entry-sides labelled nucleosome (*N* = 162), in presence of 1mM ATP.

To further study any possible role of DNA unwrapping on the entry side, we then performed a kinetic analysis of t_i_, t_du_, and t_dc_ for the entry side. t_i_ was clearly shorter on the entry side than the exit side (Fig. S16D), indicating that the unwrapping events of DNA on the entry and exit sides are unsynchronized. In contrast to what we observed with the exit side, the distributions of the t_du_ and t_dc_ data for the entry side were similar (Fig. 5E). In contrast to the marked difference in the DNA unwrapping dynamics on the entry and exit sides, the time delay between ATP injection and the first translocation event (t_r_ in Fig. 5B) was similar for both the exit and entry sides (Fig. 5F). This observation is not consistent with the recent report that time delays of hundreds second exist between the remodeling events on exit and entry sides (40). Considering all these observations, DNA unwrapping on the entry side is not strongly coupled with nucleosome translocation, and the role of DNA unwrapping on the entry side in Chd1p-mediated nucleosome remodeling is not clear.

### DNA unwrapping on the exit side dictates the remodeling speed

Our data suggest DNA unwrapping on the exit side is a critical step for a nucleosome remodeling by Chd1p. It is shown that DNA flexibility is a determining factor for DNA unwrapping/wrapping dynamics (40). The nucleosome remodeling by Chd1p is strongly affected by DNA sequences (41). We therefore studied the correlation between DNA unwrapping on the exit side and a nucleosome remodeling by Chd1p using a series of DNA sequences with variation on the exit side (Fig. S18). t_i_ and t_r_ sensitively depended on DNA sequences (Fig. 6A-B) with a strong correlation between them (Pearson’s r: 0.973, Fig. 6C). These data implicate that both the nucleosome remodeling efficiency and kinetics may be controlled by DNA sequences.

**Fig 6.**
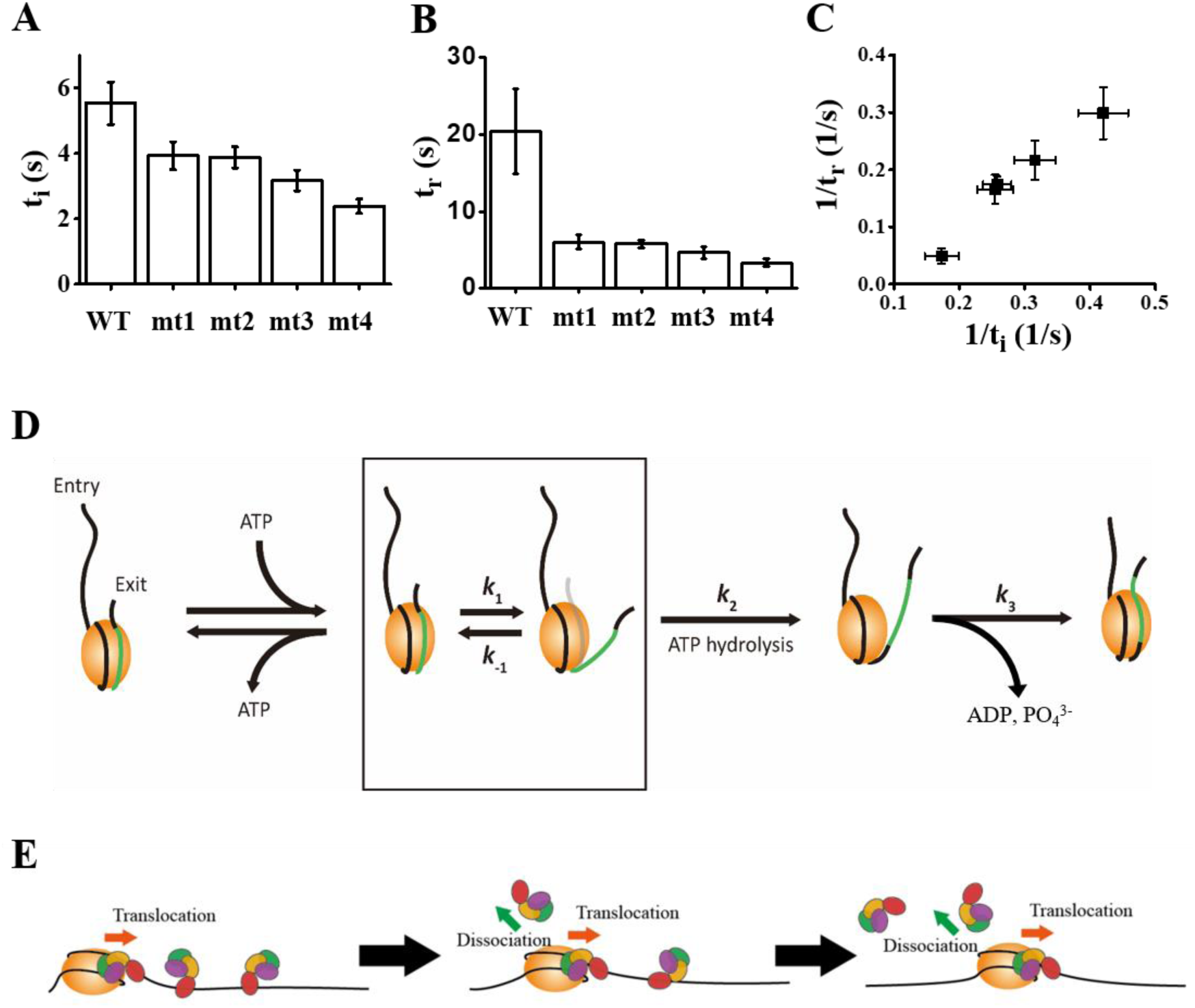
Correlation between DNA unwrapping on the exit side and nucleosome remodeling. t_i_ (A) of a series of nucleosomes with sequence variation on the exit side (Fig. S18), and their t_r_ (B). The experiments were performed in the presence of 1mM ATP. (C) Correlation plot of 1/t_r_ and 1/t_i_. The correlation coefficient (Pearson’s r) was 0.974. (D) A proposed model of nucleosome remodeling by Chd1p. (E) A proposed model of sequential and unidirectional nucleosome remodeling by multiple Chd1ps.

## DISCUSSION

Recent studies commonly suggested that superfamily 2 (SF2) helicases including several chromatin remodelers translocate on DNA with 1-bp step in general (18, 19, 42, 43). Based on structural information but without direct evidence, Chd1p was also assumed to translocate a nucleosome with 1-bp step (36). Interestingly, however, we revealed that Chd1p translocates nucleosomes in multiple base pair steps per one hydrolyzed molecule of ATP. As the ISWI and RSC remodelers were found to translocate nucleosomes in 1–2-bp steps (18, 19), larger translocation step sizes seem to be a defining characteristic of Chd1p. Without detailed information on structural differences between distinct chromatin remodelers at present, however, it is hard to present a model to explain this extraordinary nucleosome remodeling behavior of Chd1p. It is probable that there may be a larger conformational change in the ATPase domain of Chd1p than those of other chromatin remodelers, or other domains of Chd1p may help. Future biophysical studies elucidating how ATP hydrolysis energy is used to translocate a nucleosome by Chd1p are definitely required. In this work we also found that DNA on the exit side is unwrapped before most of remodeling events. Large step sizes require an increased energy cost for remodeling. DNA unwrapping might be involved in reducing the energy cost of, and thus facilitate the remodeling process.

Several static cryo-EM Chd1p-nucleosome structures and FRET experiments revealed that DNA is detached in an ATP-dependent fashion (11, 20, 36, 44), but it has remained unclear exactly what role DNA unwrapping plays during nucleosome remodeling. We revealed that DNA unwrapping on the exit side is strongly coupled with, and accelerates DNA translocation along a histone octamer. Consistent with this, the cryo-EM structures of Chd1p reported (20, 36, 45) interact with an extra DNA at the exit site where the DNA is detached from nucleosome. However, we cannot totally exclude a possibility that Chd1p also interacts the entry side of DNA when there are extra DNA at both sides. Regardless of Chd1p binding of the entry side, our data clearly show that the unwrapping at entry side is not correlated with sliding.

Although further studies are required, it is reasonably conceivable that DNA unwrapping on the exit side facilitates DNA translocation by weakening DNA-histone interactions. Although other groups have published single-molecule FRET studies of other chromatin remodelers including ACF, ISWI, and RSC (17-19), dynamic DNA unwrapping on the exit side strongly coupled with DNA translocation has never been reported. DNA unwrapping may represent another unique characteristic of Chd1p. Several previous studies suggested the existence of DNA looping during the remodeling process (46-48), but the DNA unwrapping we observed does not seem to be related to the DNA looping. In the looping model, DNA unwrapping coupled with remodeling should occur on the entry side not the exit side.

It has always been a fascinating question how molecular motors couple each step of ATP hydrolysis to their function. We revealed that ATP binding to Chd1p induces DNA unwrapping on the exit side of nucleosome, that the step of ATP hydrolysis is coupled to DNA translocation, and that the phosphate and hydrolyzed nucleotide must be released before DNA rewrapping can occur following nucleosome translocation (Fig. 6D). Therefore, it seems that the individual steps of the ATP hydrolysis cycle (i.e., ATP binding, ATP hydrolysis, and release of ADP and a phosphate) are each coupled to a distinct step of the nucleosome remodeling process (i.e., DNA unwrapping, DNA translocation, and DNA rewrapping).

It is known that Chd1p generates regularly spaced nucleosomal arrays *in vivo* (49), and, consistently with the observation, moves histone octamers to the center of short DNA fragments *in vitro* (50-53), but its exact mechanism remains unknown. We revealed that multiple Chd1ps binding to a single nucleosome sequentially move a histone octamer with a preference to the center of DNA fragments (Fig. 6E). Such a demanding task of switching Chd1ps acting on a single nucleosome can be achieved only if the interaction of Chd1p and a nucleosome is highly dynamic (21). Further single-molecule FRET studies with labeled Chd1p will elucidate the dynamic nature of the Chd1p-nucleosome interactions.

The results of several studies have suggested the different families of ATP-dependent chromatin remodelers each use a distinct nucleosome remodeling mechanism (54). This seems consistent with the presence of distinct protein domains in the ATPase subunits of each family and with the presence of distinct subunits in each remodeler complex. Consistently with this expectation, our single-molecule FRET analysis of the yeast CHD1 ATP-dependent chromatin remodeler revealed that the mechanism by which Chd1p remodels nucleosomes is distinct from those of remodelers from the ISWI and SWI/SNF families. It would be interesting to examine whether other remodelers in CHD family have similar properties to Chd1p. Considering that Chd1p, ISWI, and SWI/SNF are all members of the SF2 superfamily (17-19), it remains unclear exactly what about Chd1p makes it function in this way. To address the exact mechanism of these characteristic features of Chd1p might require a series of high-resolution structures in different ATP hydrolysis states.

## METHODS

### Protein expression, and purification

Yeast Chd1 cDNA was cloned into pFastBac vector for expression in Sf9 insect cells. N-terminal His-tagged Chd1 was transduced in Sf9 via baculovirus system. Transduced cells were cultured for 45 h, harvested and resuspended in a buffer containing 300 mM NaCl, 5% Glycerol, 50 mM Tris pH 8.0, and cOmplete™, Mini, EDTA-free Protease Inhibitor Cocktail (Roche). His-tagged Chd1 was purified by Ni-NTA agarose affinity chromatography (Qiagen) and treated with homemade TEV protease to remove the tag, followed by ion exchange and size exclusion chromatography (GE Healthcare) in a buffer containing 300mM NaCl, 50mM Tris pH 8.0, 1mM DTT and 20% Glycerol. The SNAP tagged Chd1p was also expressed in Sf9 insect cells and purified as the wild-type Chd1p. *Xenopus laevis* histones including H2A T120C were cloned into pET vector and expressed in *E. coli* (BL21 DE3 pLyss) as an insoluble form. Each histone in inclusion body was unfolded in a buffer containing 6 M guanidine HCl, 20 mM Na-Ac pH 5.2, and 5 mM DTT and purified by size exclusion and ion exchange chromatography (GE Healthcare). Purified histones were lyophilized and resolved with the same molecular ratio in the buffer used for unfolding process. Resolved histones were mixed with each other and dialyzed in a buffer containing 2 M NaCl, 10 mM Tris pH 7.5, 1 mM EDTA, and 5 mM β-mercaptoethanol to be assembled into histone octamer.

### DNA and nucleosome preparation

The standard nucleosomes used in this work were designed to have a 78-bp spacer on the entry side and a 3-bp spacer on the exit side (Figure S1A). DNA fragments containing 601 nucleosome positioning sequence (37) were generated by PCR as described (33). Primers with Cy5 at the 5’ end, internal amine modification, or biotin were purchased from IDT. The amine modified primers were labeled with Cy5 NHS ester. PCR products were purified by 5% native PAGE gel. Nucleosomes were reconstituted with histone octamer containing either Cy3-labeled H2A T120C or Cy3-labeled H4 R45C and Cy5-labeled DNA fragment using salt gradient dialysis. To immobilize the SNAP tagged Chd1p described above, the biotin-Chd1p was prepared by mixing the SNAP-Chd1p and SNAP-biotin® (New England BioLabs) in a 1:2 molar ratio and incubated at room temperature for 30min. Then, biotin-Chd1p were immobilized on a PEGylated surface using biotin-streptavidin conjugation like above and incubated with Nucleosome substrates without biotin for 2-3 minutes.

### Single-molecule FRET experiment

Quartz slides and coverslips were cleaned with piranha solution (mixture of 3:1 concentrated sulfuric acid:30% [v/v] hydrogen peroxide solution), coated with aminopropyl silane first, and then with mixture of PEG (m-PEG-5000, Laysan Bio) and biotin-PEG (biotin-PEG-5000, Laysan Bio). A flow cell was assembled by combining a coverslip and a quartz slide with double-sided adhesive tape (3M). For convenient buffer exchange, polyethylene tubes (PE50; Becton Dickinson) were connected to the flow cell. Nucleosome substrates were immobilized on a PEGylated surface using biotin–streptavidin conjugation and then incubated with Chd1p(180 nM) for 2-3 minutes. Single-molecule FRET experiments were performed in an imaging buffer (10 mM Tris–HCl pH 8.0, 100 mM KCl, 3 mM MgCl_2_, 5 mM NaCl) containing a gloxy oxygen scavenging system (a mixture of glucose oxidase (1 mg/ml, Sigma), catalase (0.04 mg/ml, Sigma), glucose (0.4% (w/v), Sigma), and Trolox (2 mM, Sigma)). During imaging, various concentration of ATP or ATP analog was injected with imaging buffer to initiate the reactions by Chd1p. For Mg^2+^ flow experiment, surface immobilized nucleosomes were incubated with 180 nM Chd1p and 100μM ATP in imaging buffer (with 10 mM EDTA instead of MgCl_2_), and then an imaging buffer containing 3mM MgCl_2_ was injected during imaging. Single-molecule fluorescence images were acquired at a frame rate of 10 Hz using a home-built prism-type total internal reflection fluorescence microscope equipped with an electron-multiplying charge coupled device camera (Ixon DV897; Andor Technology). Cy3 and Cy5 were alternately excited with 532nm and 633nm lasers using the ALEX (Alternative Laser EXcitation) technique (35). Experimental temperature was maintained at 30°C using a temperature control system (Live Cell Instruments). Data acquisition and FRET trace extraction were done using homemade programs written in LabView (National Instruments) and IDL (ITT), respectively. The intensity traces of Cy3 and Cy5 were analyzed using a custom made Matlab (MathWorks) script. The FRET efficiencies were calculated from the donor (ID) and acceptor (IA) fluorescence intensities as EFRET = IA/(IA + ID).

## ACKNOWLEDGMNETS

This work was supported by a grant (NRF-2019R1A2C2005209) to SH, and by grants (NRF-2016R1A2B3006293, NRF-2016K1A1A2912057) to J.S. from National Research Foundation of Korea.

## AUTHOR CONTRIBUTIONS

JK, JYL, CK, and YL performed the experiments, and analyzed data. All authors contributed to writing of the paper.

## Supplementary Information

**Figure S1.**
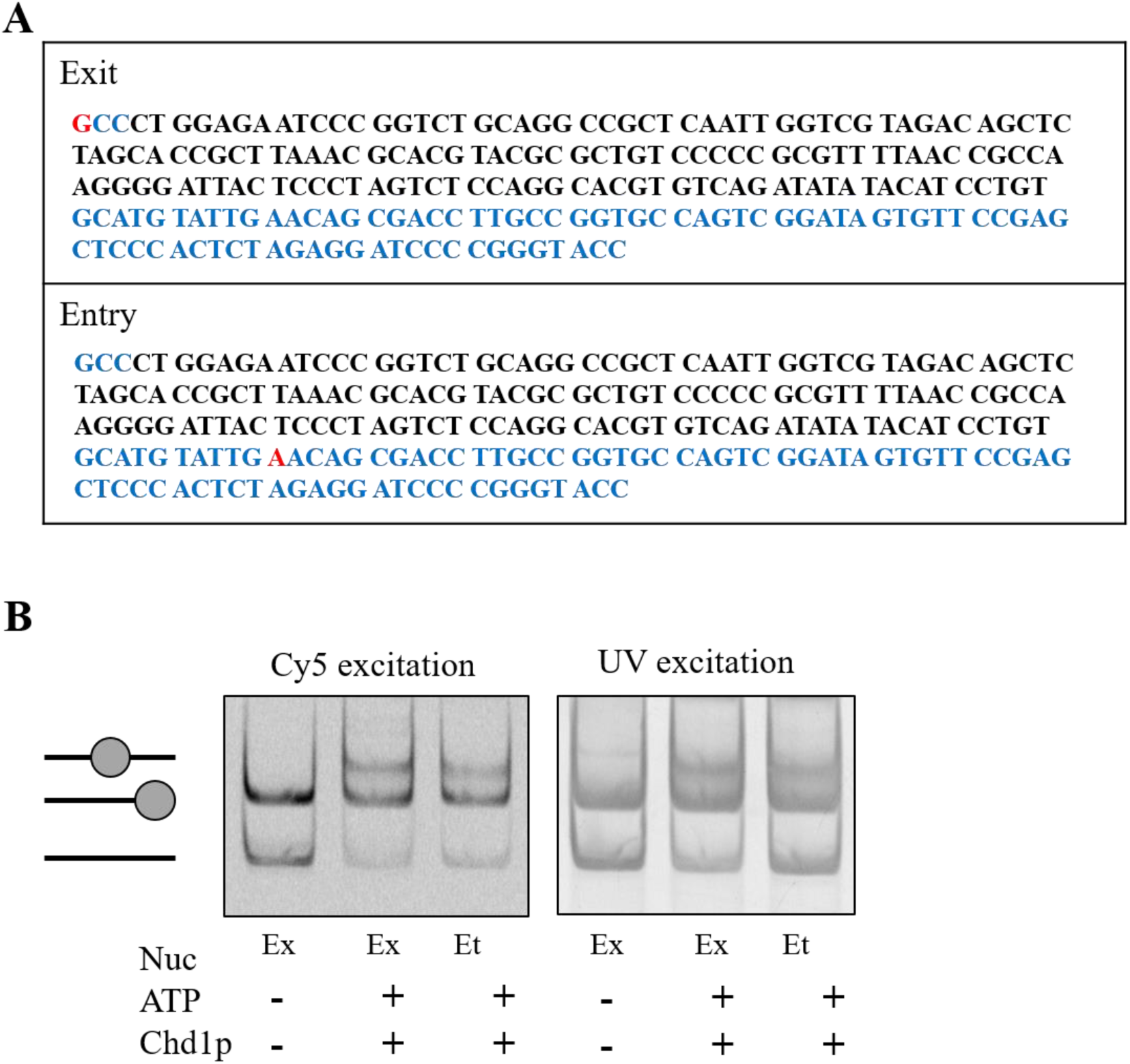
DNA sequences and remodeling activity of Chd1p on labelled nucleosomes. (A) DNA sequences used in the study. The DNA fragments consist of 601 nucleosome positioning sequence (black) with the 3 bp linker on the exit side and the 78 bp on the entry side (blue). The Cy5 labeling position is marked in red. (B) Remodeling activity of Chd1p on fluorophore-labeled nucleosomes. Bulk remodeling assays were carried out in 10 mM Tris-HCl pH 8.0 with 100 mM KCl, 3 mM MgCl_2_, 5 mM NaCl, 2.5 mM ATP, 700 nM of Chd1p, and 280 nM nucleosomes at 30°C for 20 min. The products were examined using a 5% native PAGE gel. (Nuc, Nucleosome; Ex, exit-side nucleosome; Et, entry-side nucleosome)

**Figure S2.**
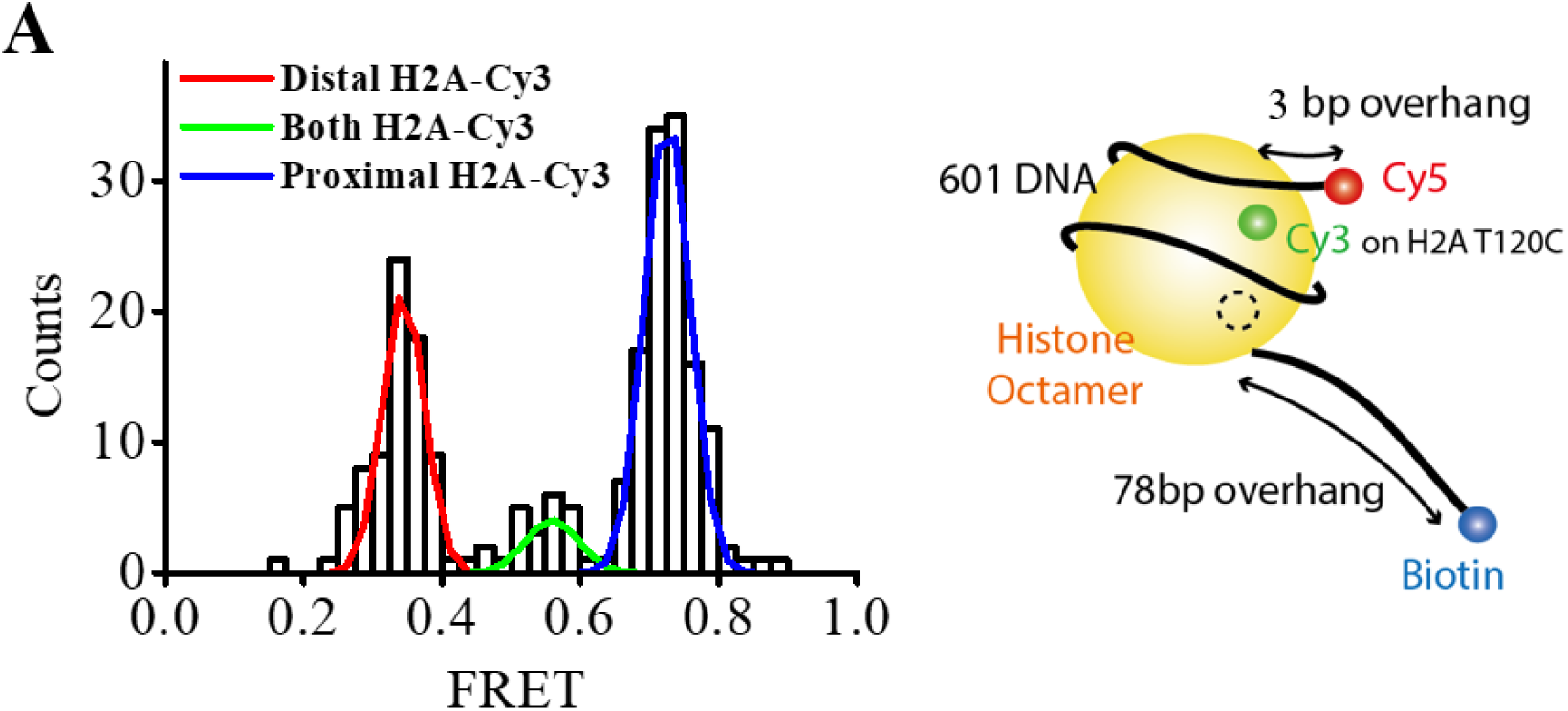
FRET histogram of the exit-labeled nucleosome. (A) FRET histogram and labeling scheme of the exit-side labelled nucleosomes. The histogram shows three FRET peaks. The low and high FRET corresponds to species labeled at the distal and proximal H2A, respectively. The mid FRET corresponds to doubly-labeled species. We selected only high FRET species for data analysis.

**Figure S3.**
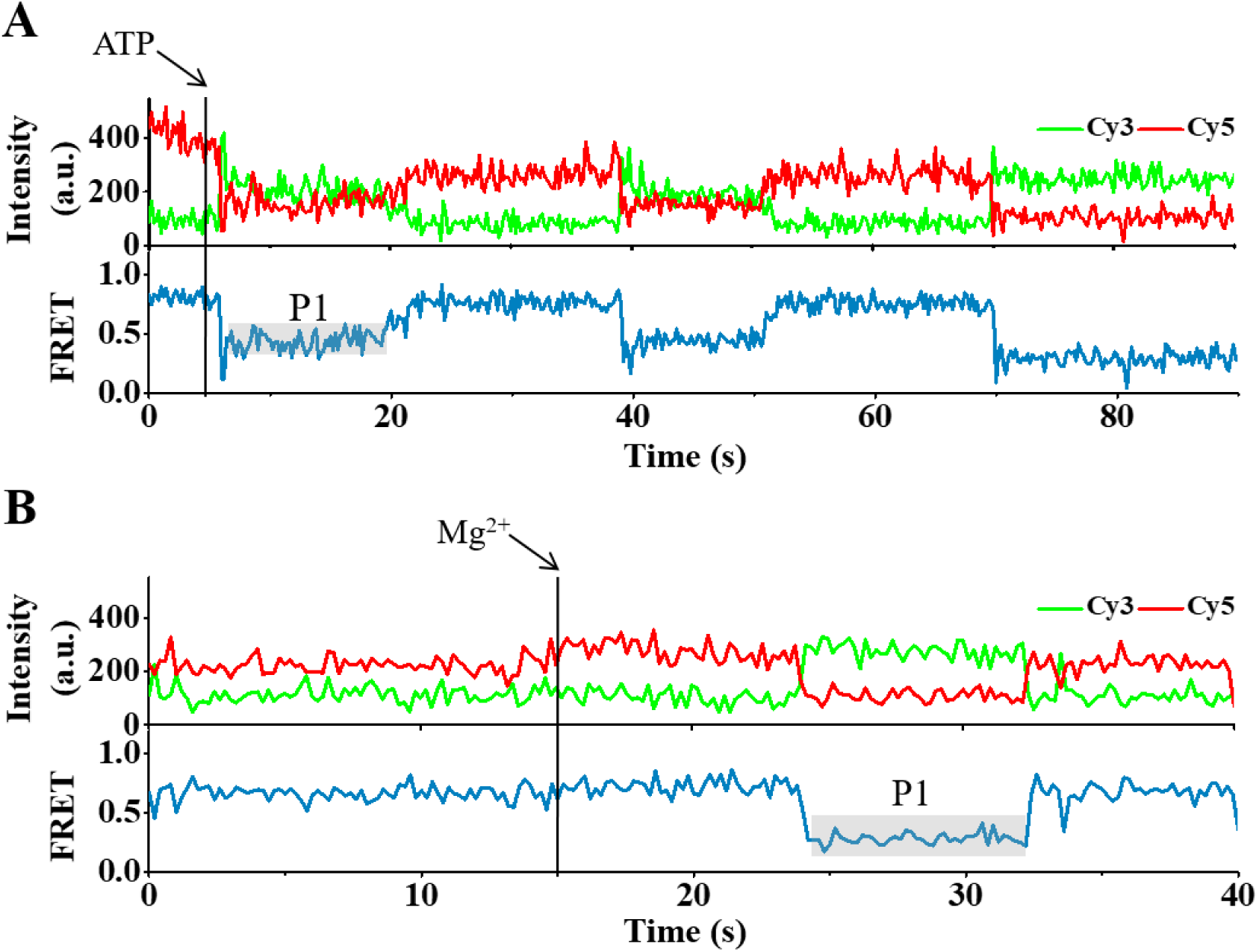
Nucleosome remodeling by Chd1p is reversible. (A) A representative fluorescence and FRET time traces showing back-and-forth translocation of nucleosome on the exit side by CHD1. The reaction was started by injecting ATP. (B) A representative fluorescence and FRET time trace showing back-and-forth translocation on the exit side. Chd1p-nucleosome complex was preincubated with ATP and EDTA, and the remodeling reaction was initiated by injecting Mg^2+^.

**Figure S4.**
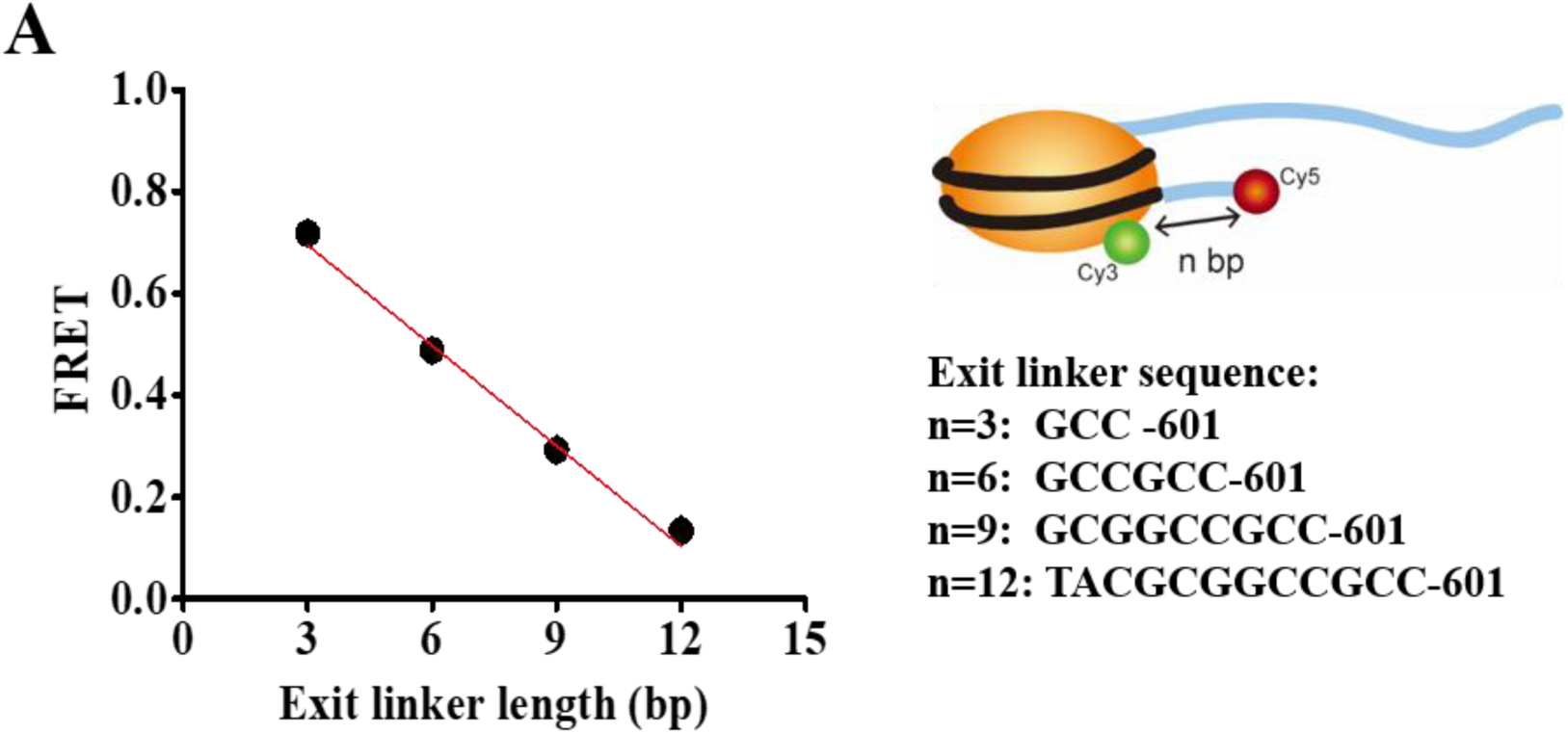
Linear relation between FRET efficiency and the exit linker length. (A) Calibration data for step size determination. FRET was obtained from nucleosomes with varying linker DNA lengths (n= 3, 6, 9, 12 bps) on the exit side. The linear-fit of the data had a slope of -0.066 ± 0.0004 per base pair.

**Figure S5.**
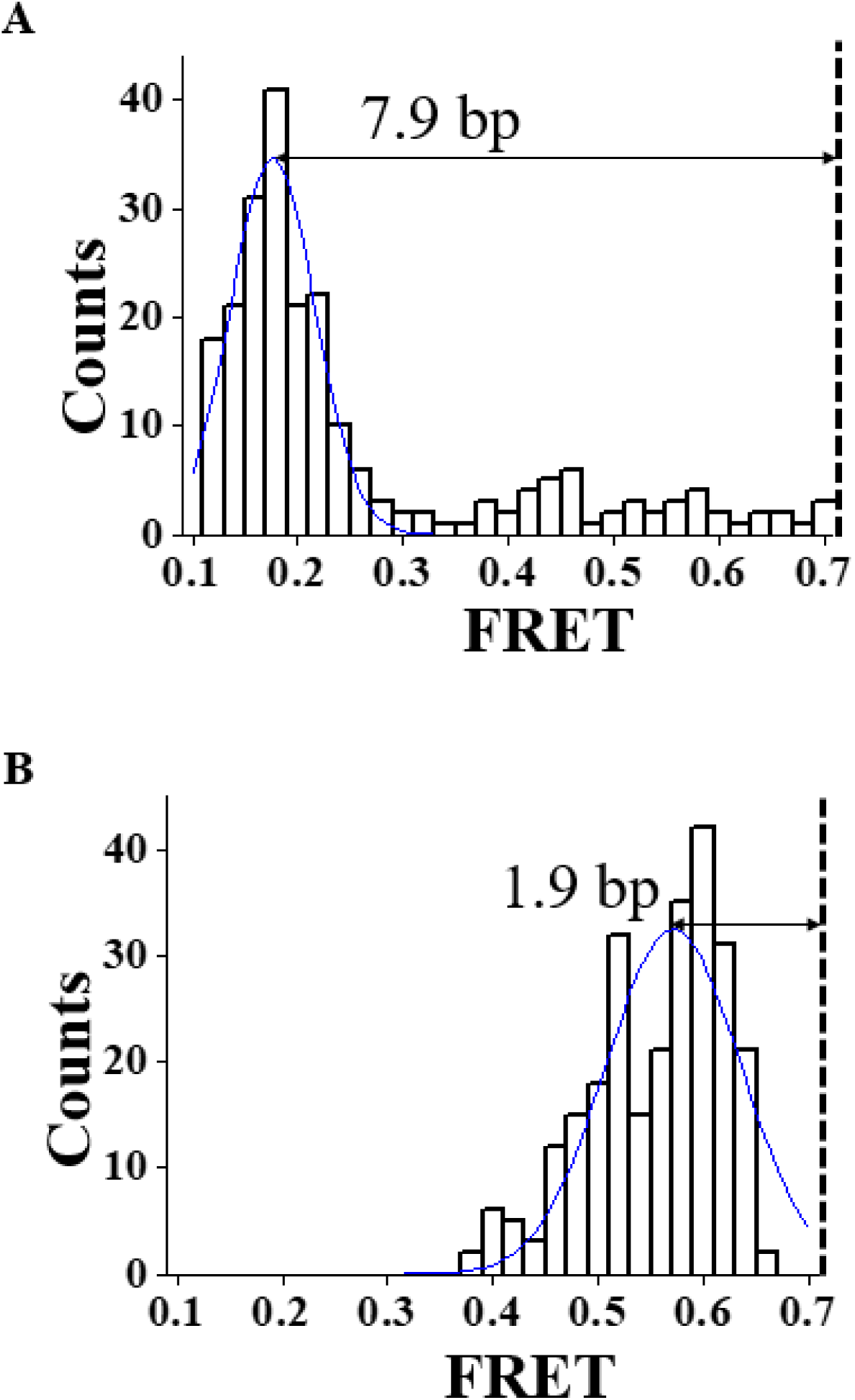
Observation of remodeling substeps of ACF complex. (A) A FRET change histogram of ACF remodeling events at 1 mM ATP concentration. (B) A FRET change histogram of ACF remodeling events at 50μM ATP and 500 μM ATPγS concentrations. The existence of remodeling substeps is clear.

**Figure S6.**
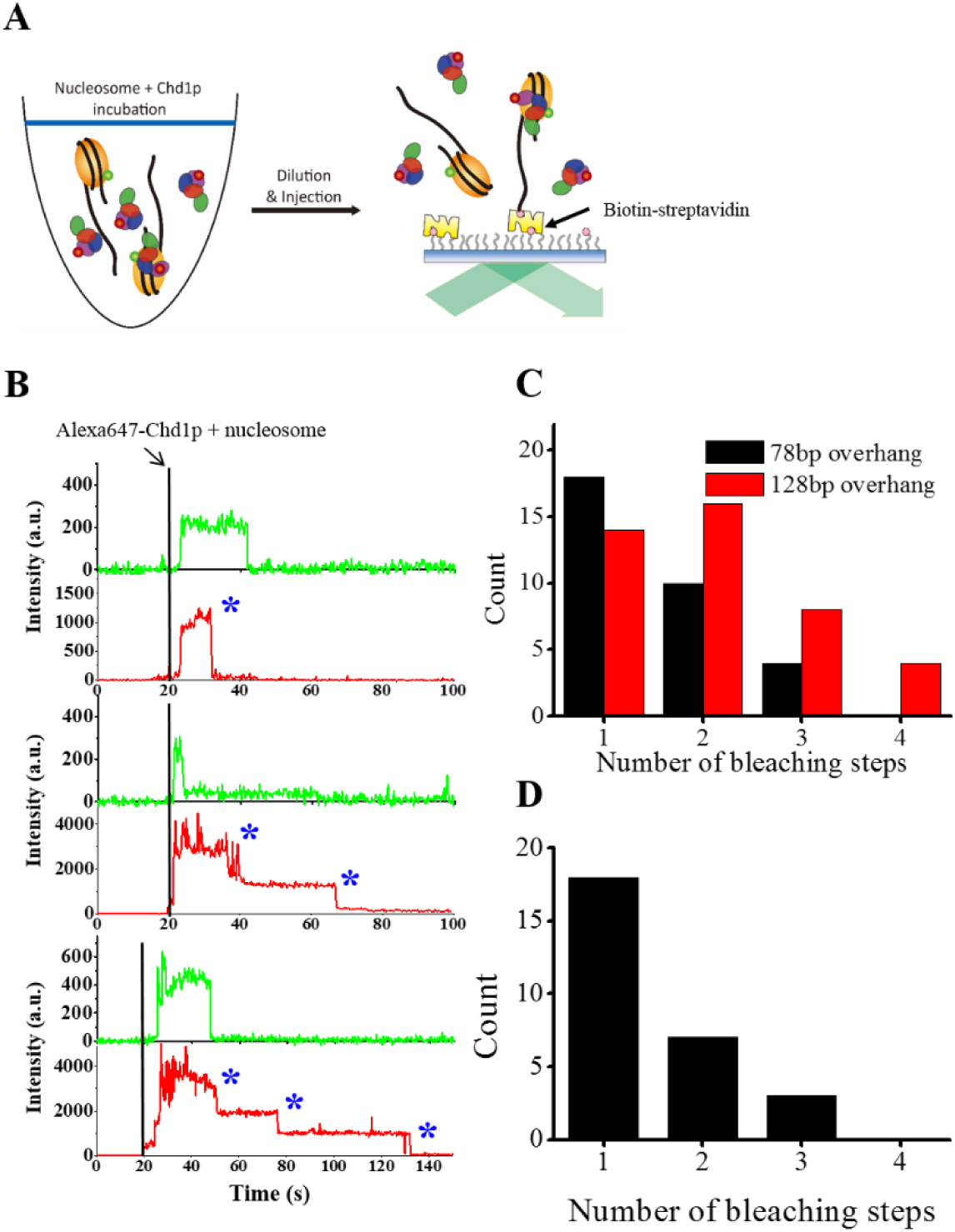
The number of Chd1p on a single nucleosome. (A) Experimental scheme for Chd1p counting. Mixture of 15nM Cy3-labeled Nucleosome and 200nM Alexa647-labeled Chd1p in T50 buffer (50mM NaCl, 10mM Tris-HCl (pH 8.0)) was incubated for 3 minutes, and 100x diluted with the imaging buffer. The diluted sample was injected into the imaging chamber while single-molecule images were being taken. Binding of a nucleosome/Chd1p complex on surface-immobilized streptavidin was monitored using single-molecule fluorescence microscope. (B) Representative fluorescence intensity time traces of Cy3 (green) and Alexa647 (red) showing the binding events (simultaneous appearance of Cy3 and Alexa647). Photobleaching steps of Alexa647 are indicated by blue asterisks. (C) Distribution of Alexa647 photobleaching step number observed in (B), using a DNA substrate with a 78bp DNA overhang, the one described in figure S1A (black), and a DNA substrate with a 128bp DNA overhang (red). Based on the distribution and the labeling efficiency of Alexa647 (80%), the fractions of a nucleosome with one, two, and three Chd1ps were estimated as 47.8%, 30.4%, and 21.7%, respectively for the case of 78bp DNA overhang sample. (D) Distribution of Alexa647 photobleaching step number observed after wash-out. A DNA substrate with a 78bp DNA overhang was used.

**Figure S7.**
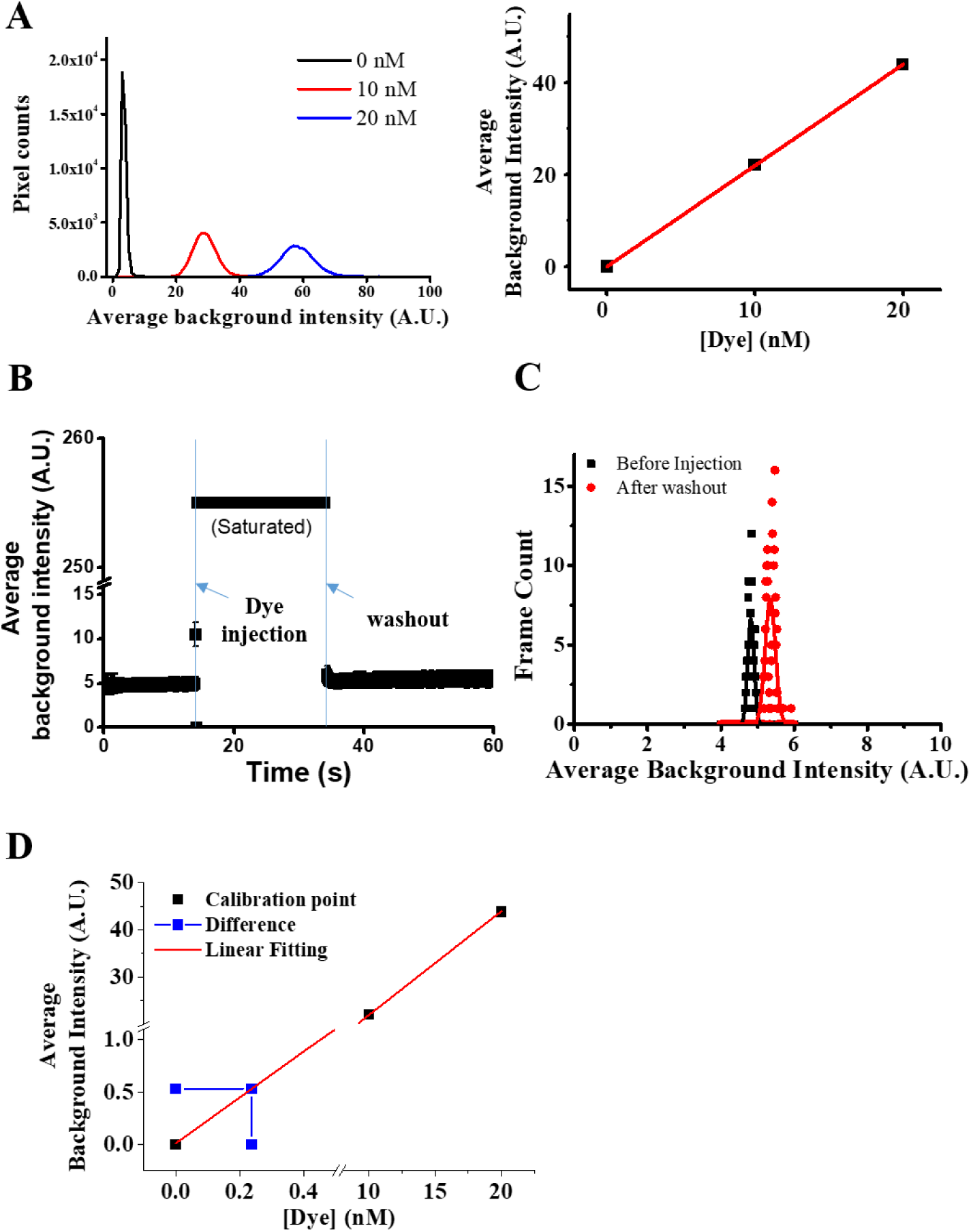
Characterization of the buffer exchange efficiency of the flow system I. (A) Histograms of EM-CCD pixel count in the presence of 0nM, 10nM, 20nM free Cy5 (left). The average background intensity and Cy5 concentration exhibited a nice linear relation (right). (B) Average background intensity before the injection and after the removal of Cy5. Cy5 (500nM) was injected into the chamber at 15s, and then the chamber was washed out with buffer containing no Cy5 (volume: 300 μl, flow rate: 100 μl/s) at 35s. (C) Background intensity distribution before the injection (black) and after washout of Cy5 (red). (D) Buffer exchange efficiency. The average background intensity after the washout was 0.53 (blue), corresponding to Cy5 concentration of 0.24 nM. Therefore, the buffer exchange system is estimated to remove 99.95% of solutes.

**Figure S8.**
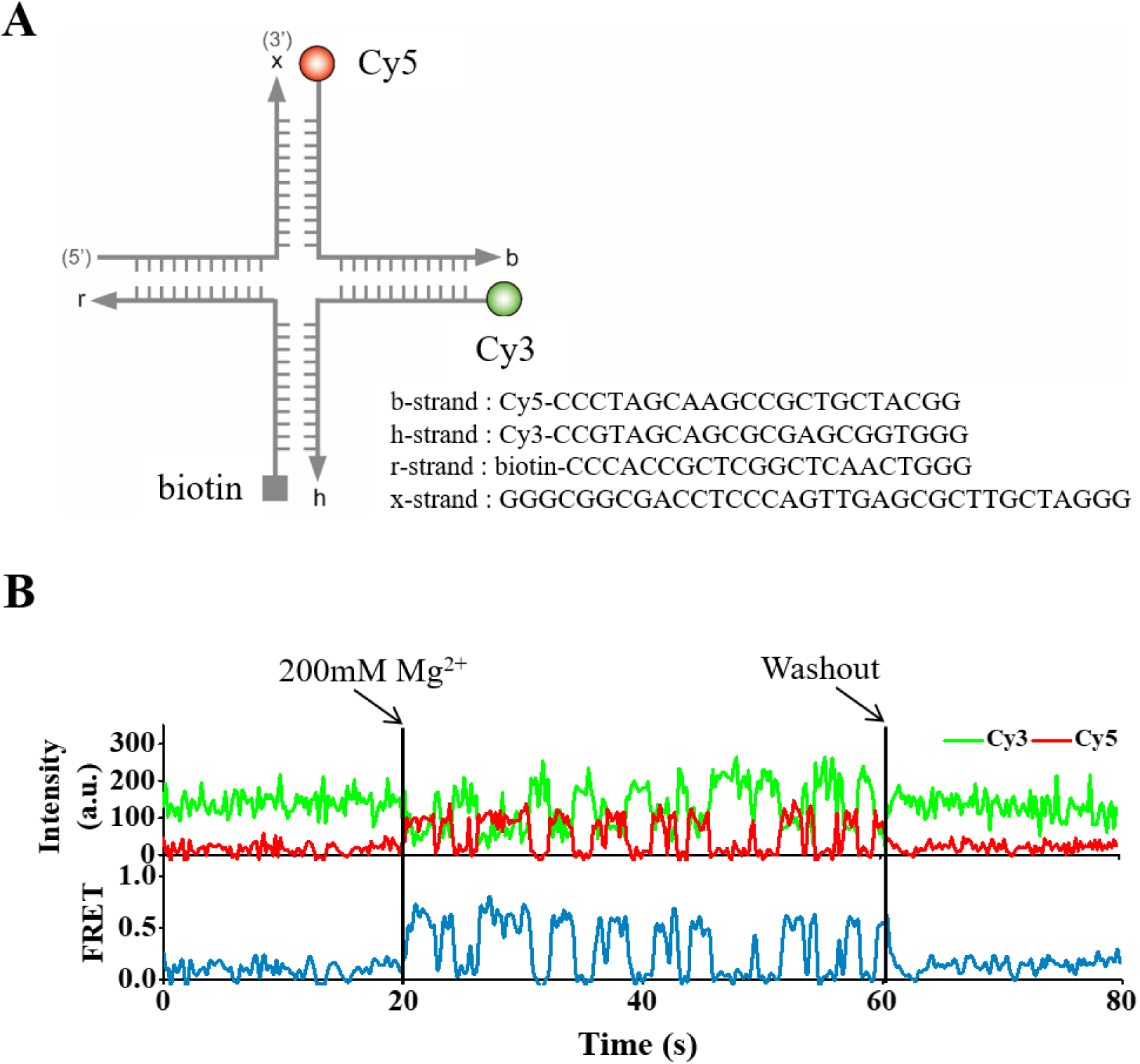
Characterization of the buffer exchange efficiency of the flow system II. (A) DNA sequences of the Holliday Junction and fluorophore labeling scheme. (B) Representative fluorescence intensity times traces during 200 mM Mg^2+^ injection and washout with 300μl of Mg^2+^-free imaging buffer. The absence of FRET dynamics after the washout indicates that Mg^2+^ remaining after the washout is negligible.

**Figure S9.**
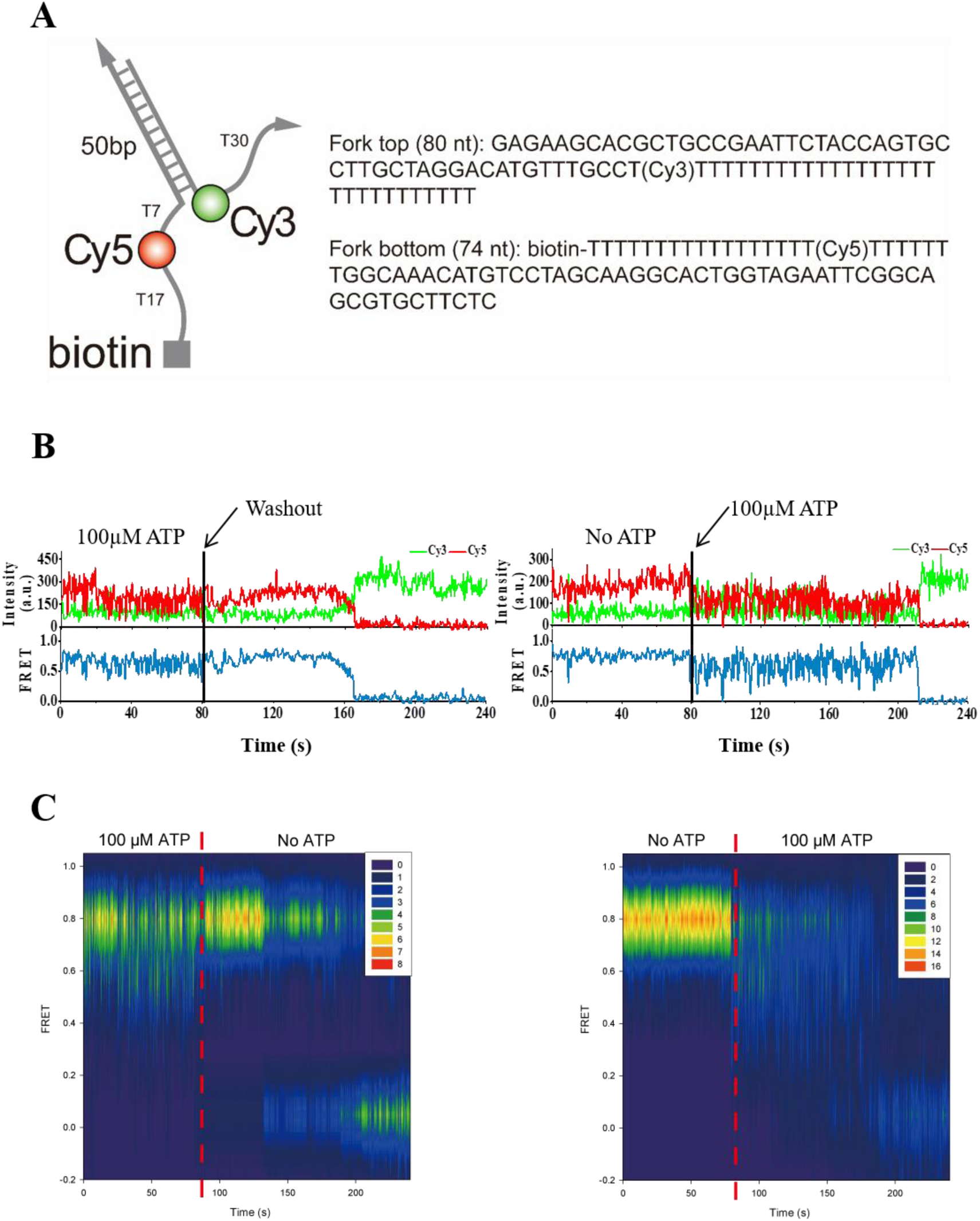
Characterization of the buffer exchange efficiency of the flow system III. (A) DNA sequences of forked DNA and fluorophore labeling scheme. (B) Representative fluorescence intensity times traces showing repetitive unwinding by HIM-6 (*C. elegans* BLM helicase ortholog) only in the presence of ATP (100 μM). (C) Contour plots of FRET efficiency of the experiments described in (B). Dashed red lines indicate buffer exchange points.

**Figure S10.**
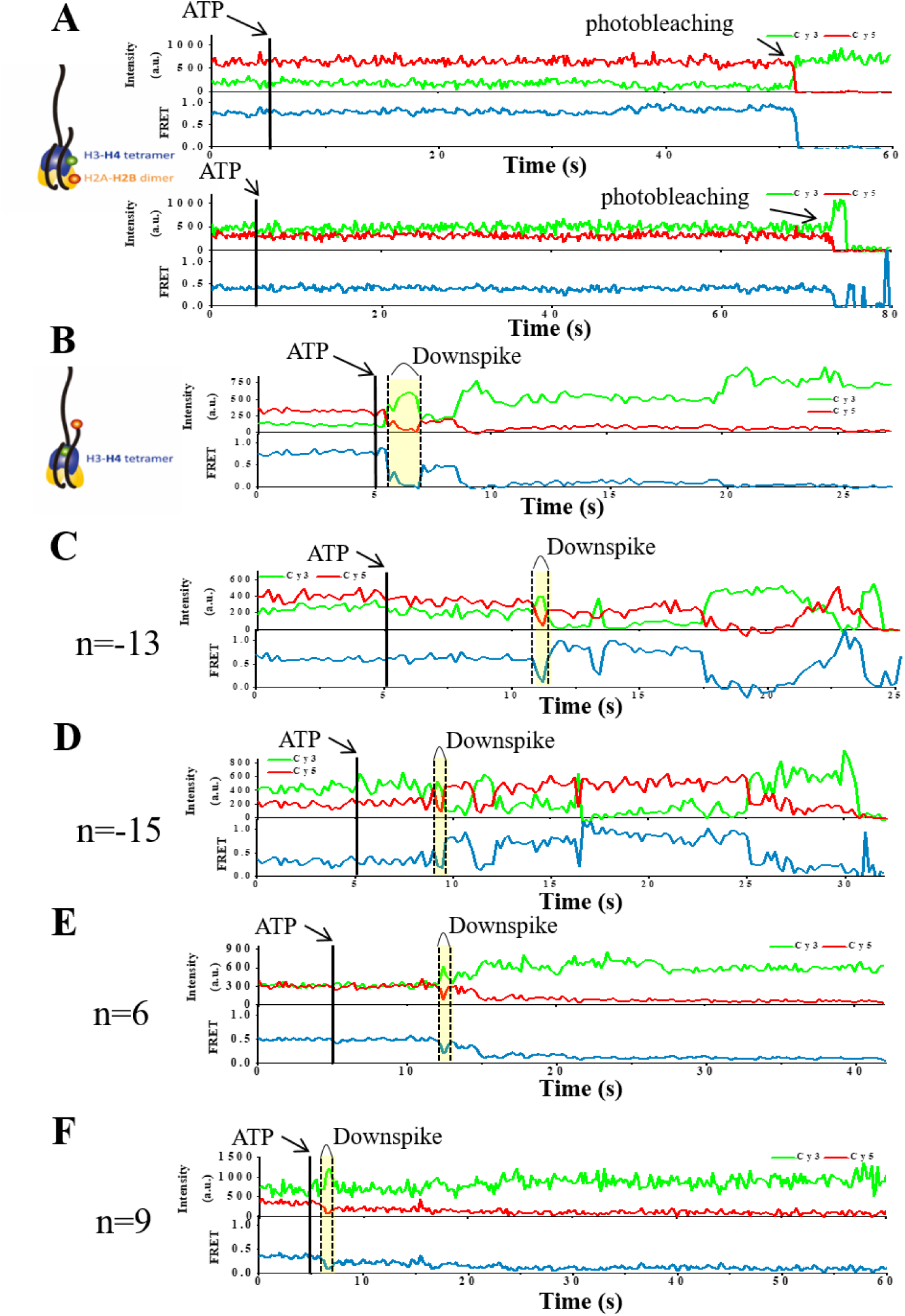
FRET down spikes on the exit side under different labeling schemes. (A) No dynamics was observed with Cy3 labeled on H4 E63C and Cy5 on H2A T120C. On the other hand, down spikes were observed with (B) Cy3 labeled on H4 R45C and Cy5 on DNA end (n=3), or (C-F) Cy3 labeled on H2A T120C, and Cy5 labeled on several different positions near the exit-side DNA. (what does the negative, positive means? Where is n=0)

**Figure S11.**
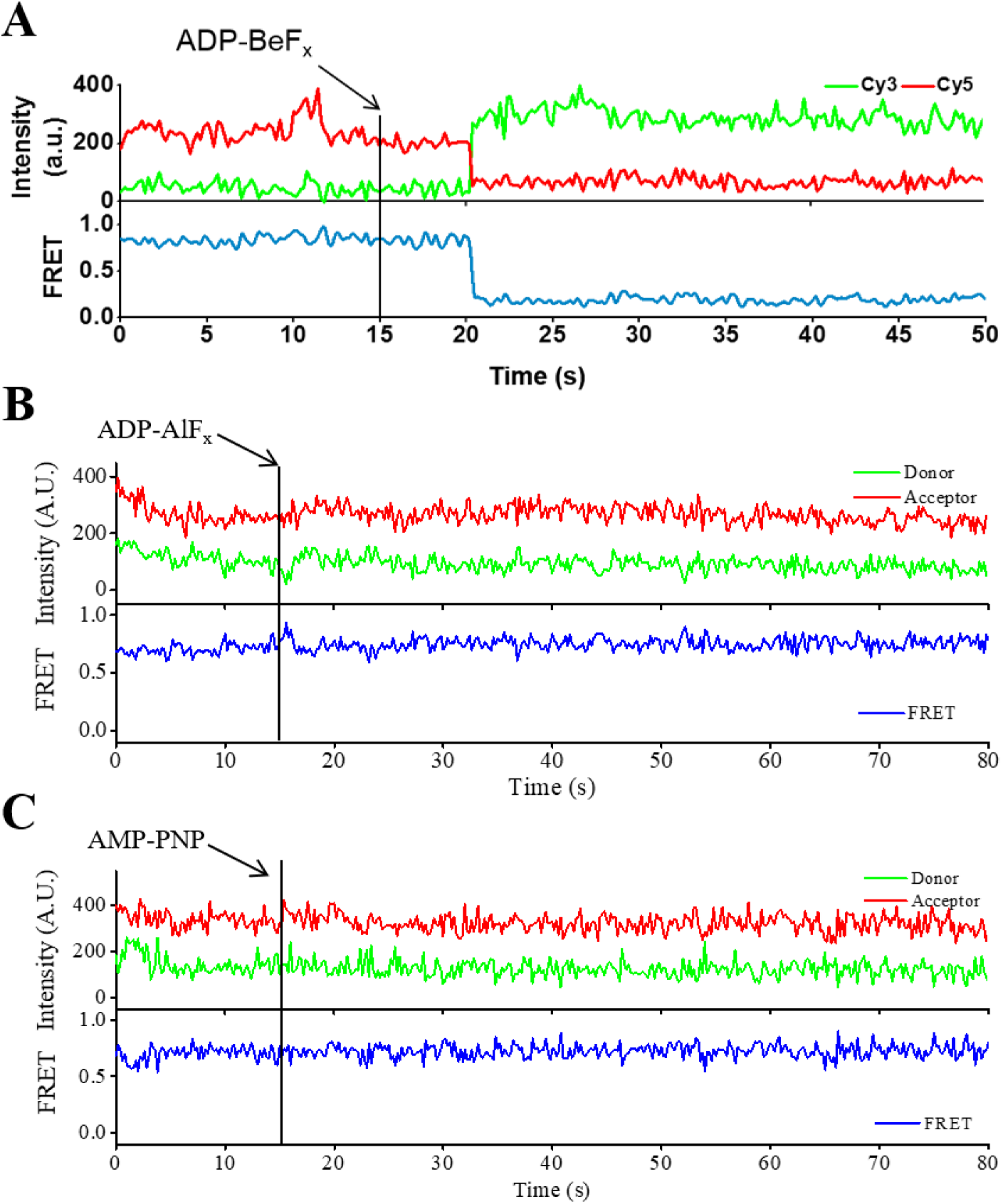
FRET down spikes on the exit side in the presence of various ATP analogs. Stable FRET down spikes were observed in the presence of (A) 1mM ADP-BeF_x_ (1mM ADP, 5mM BeCl_2_, 25mM NaF), but no down spike was observed in the presence of (B) 1mM ADP-AlF_x_ (1mM ADP, 5mM AlCl_3_, 25mM NaF), or (C) 1mM AMP-PNP.

**Figure S12.**
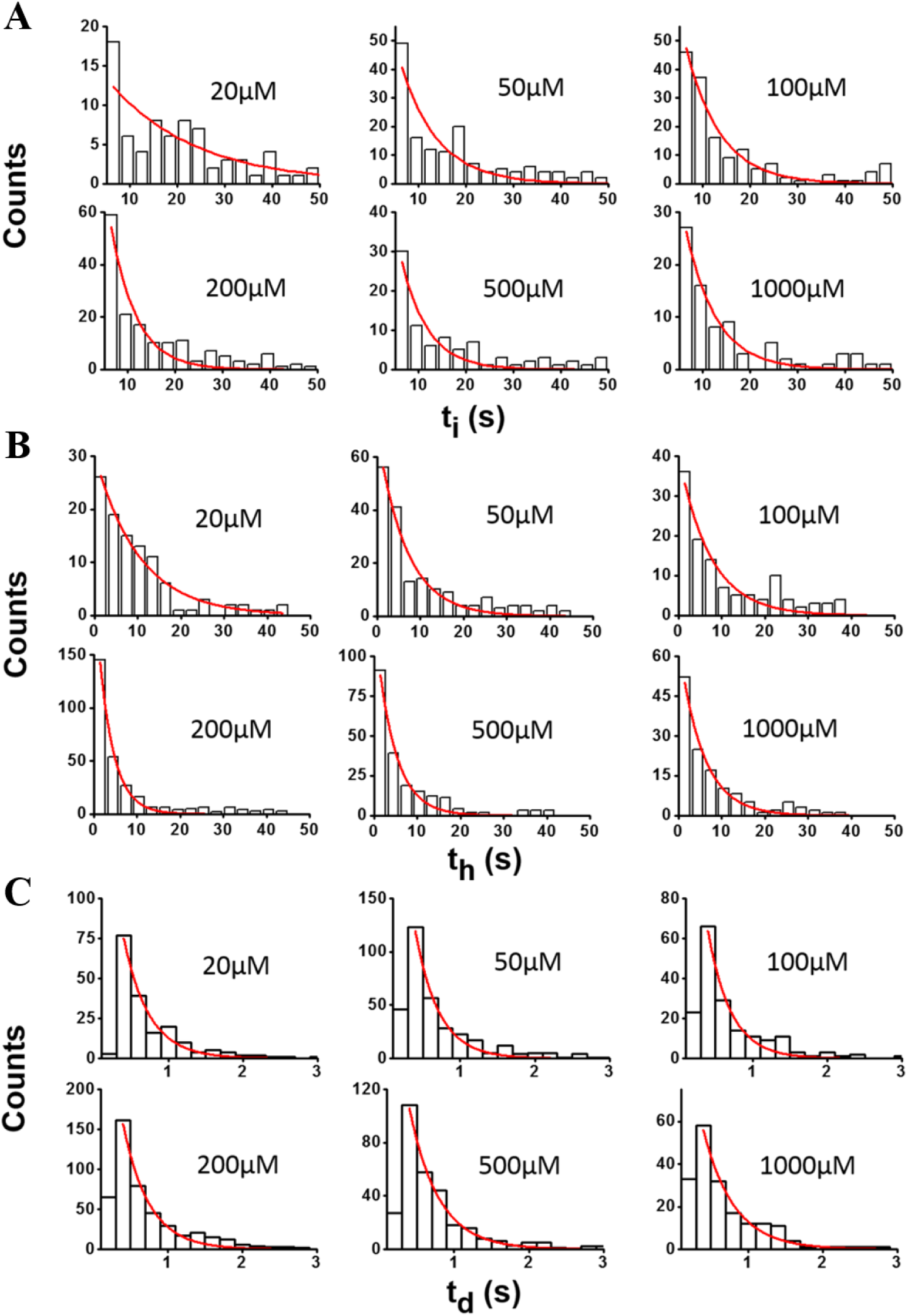
ATPγS titration of kinetic parameters of the exit side unwrapping. ATPγS titration of (A) t_i_, (B) t_h_, and (C) t_d_ of the exit side nucleosome, and their fit to single-exponential functions (red lines).

**Figure S13.**
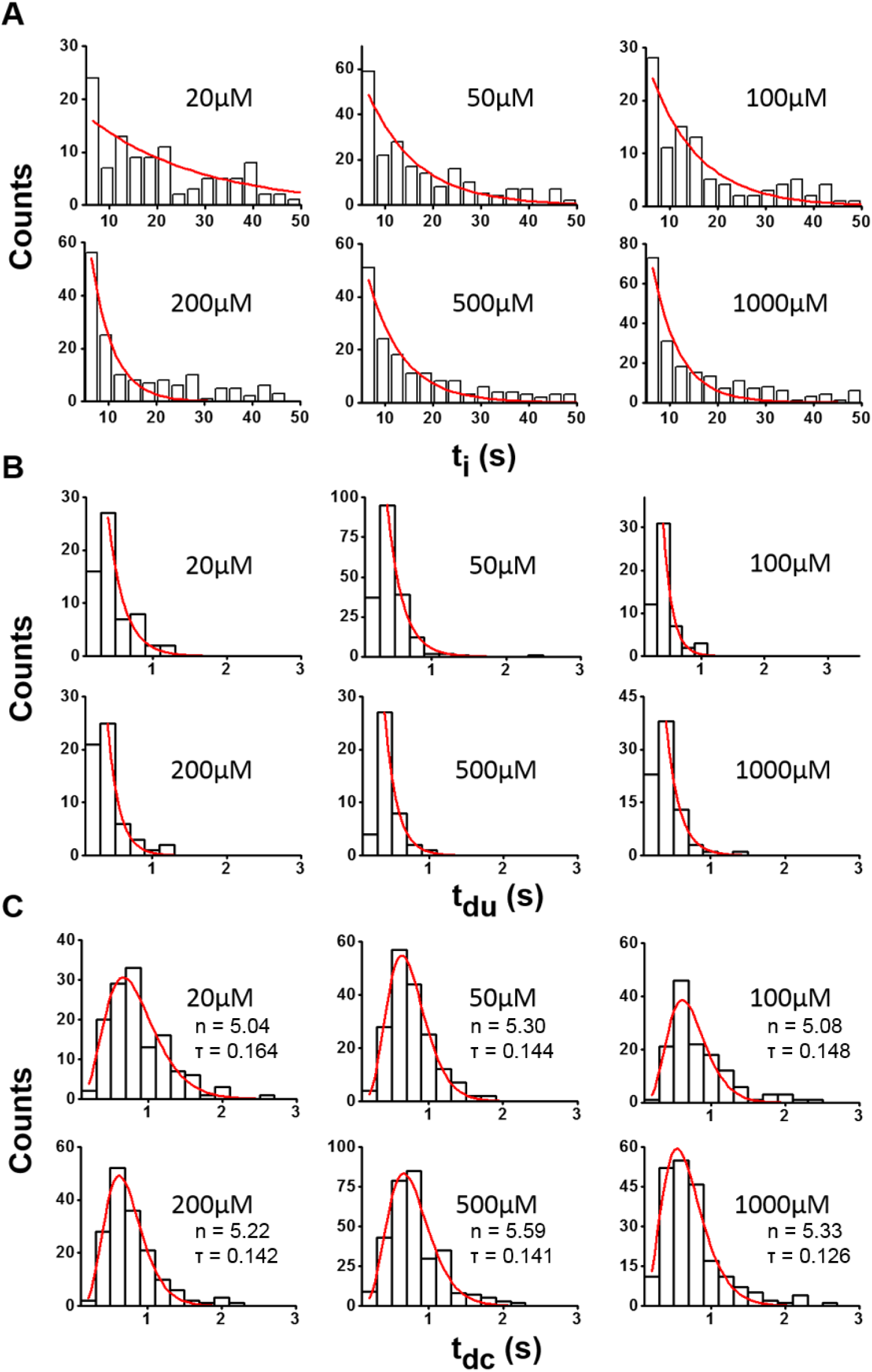
ATP titration of kinetic parameters of the exit side remodeling. ATP titration of (A) t_i_, (B) t_du_, and (C) t_dc_ of the exit side nucleosome. t_i_ and t_du_ were fitted to single-exponential functions (red lines). t_dc_ was fitted to gamma functions.

**Figure S14.**
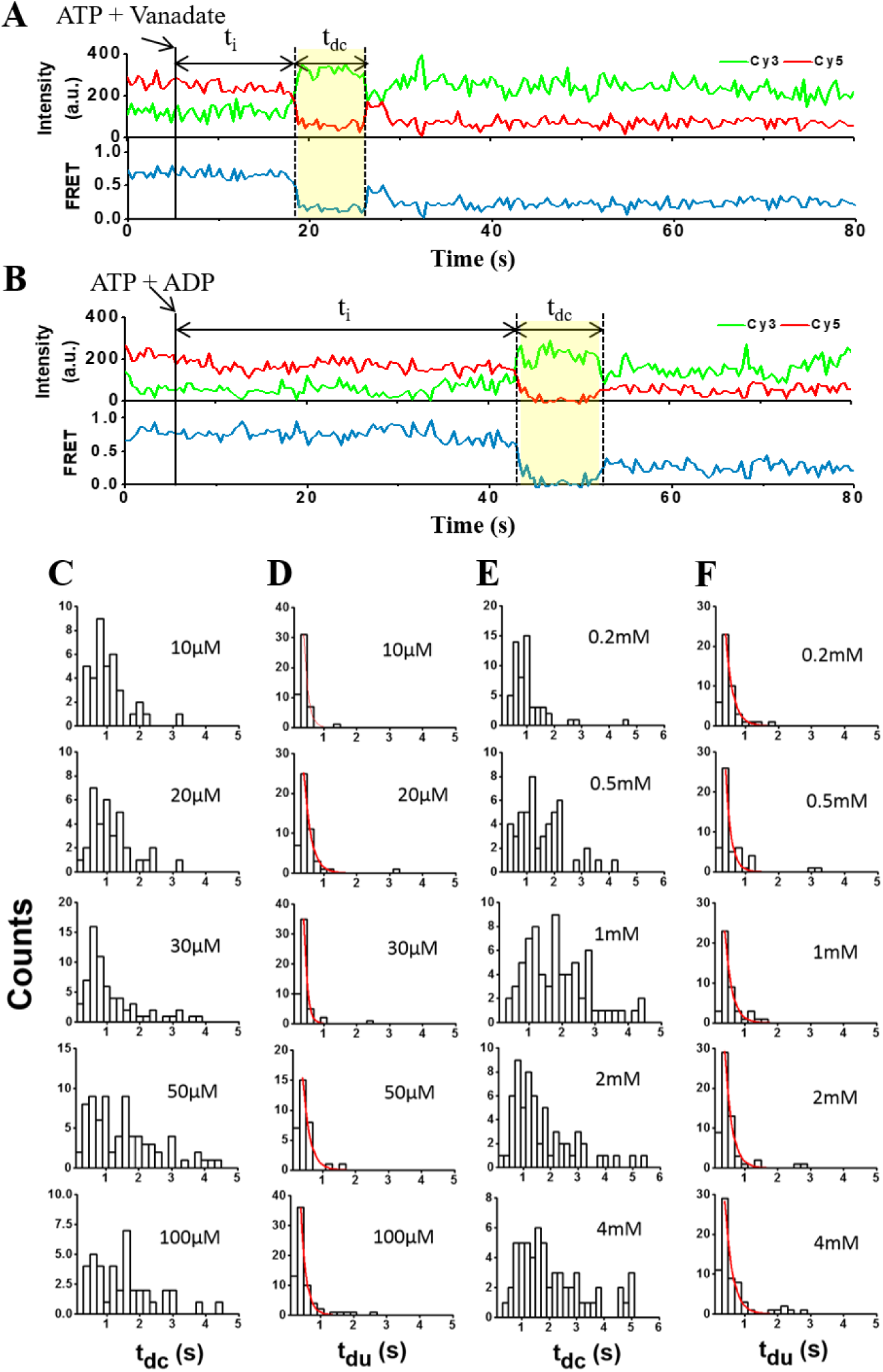
Effects of vanadate and ADP on remodeling kinetics. Representative fluorescence intensity time traces showing remodeling events in the presence of 1mM ATP and 100μM vanadate (A), or in the presence of 100mM ATP and 4mM ADP (B). Vanadate titration of (C) t_dc_, and (D) t_du_ for the nucleosome exit side at 1 mM ATP. t_du_ histograms were fitted to single-exponential functions (red lines). ADP titration of (E) t_dc_, and (F) t_du_ for the nucleosome exit side at 100 μM ATP. t_du_ histograms were fitted to single-exponential functions (red lines).

**Figure S15.**
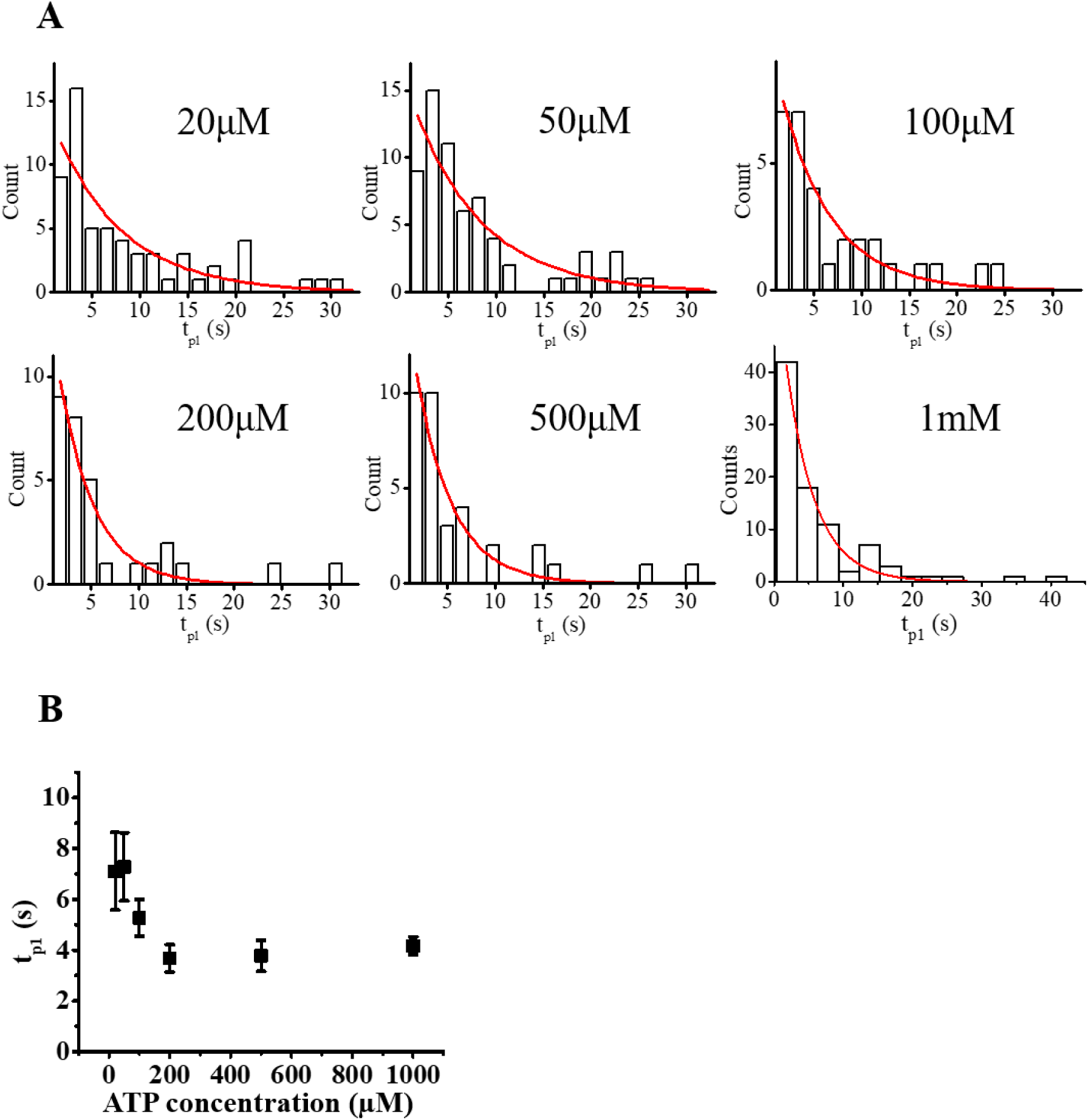
ATP dependence of t_p1_. (A) Histograms of t_p1_ at varying ATP concentration. (B) ATP titration of t_p1_. The data were obtained by fitting histograms to single-exponential functions (red lines in (A)).

**Figure S16.**
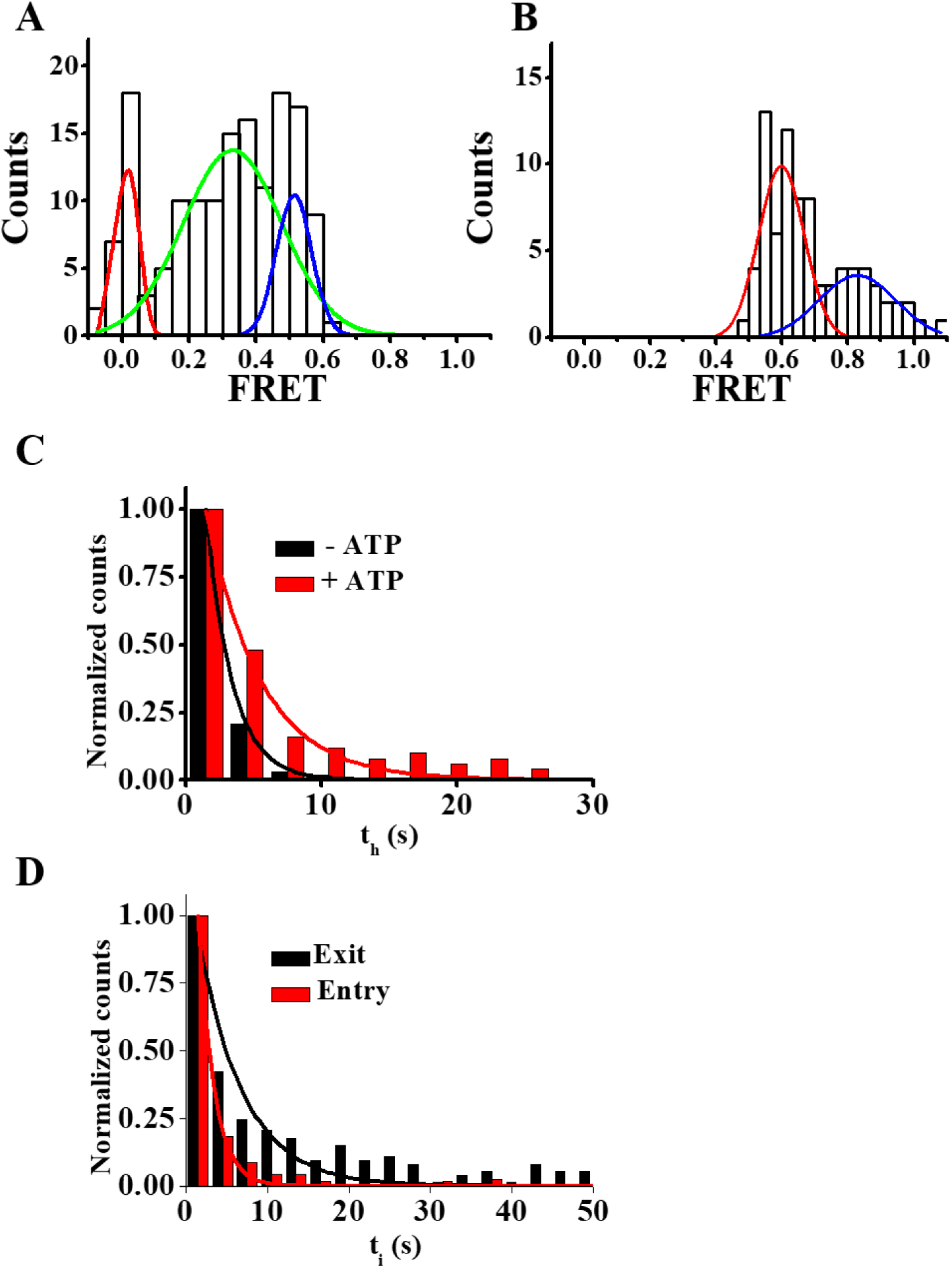
Supplementary data to characterize the entry-side remodeling. (A) FRET histogram of the entry-side nucleosome before remodeling. Like the exit-side nucleosome, three FRET peaks were observed. Only the high FRET population was used for further analysis. (B) FRET histograms after first remodeling. As with the exit-side nucleosome, two distinct FRET distributions were observed. (C) High FRET dwell time (t_h_) on the entry side with (red) and without ATP (black), and their fit to single-exponential functions (solid lines). (D) Comparison of the initiation time (t_i_) of the entry-side (red) and the exit-side nucleosome (black), and their fit to single-exponential functions (solid lines).

**Figure S17.**
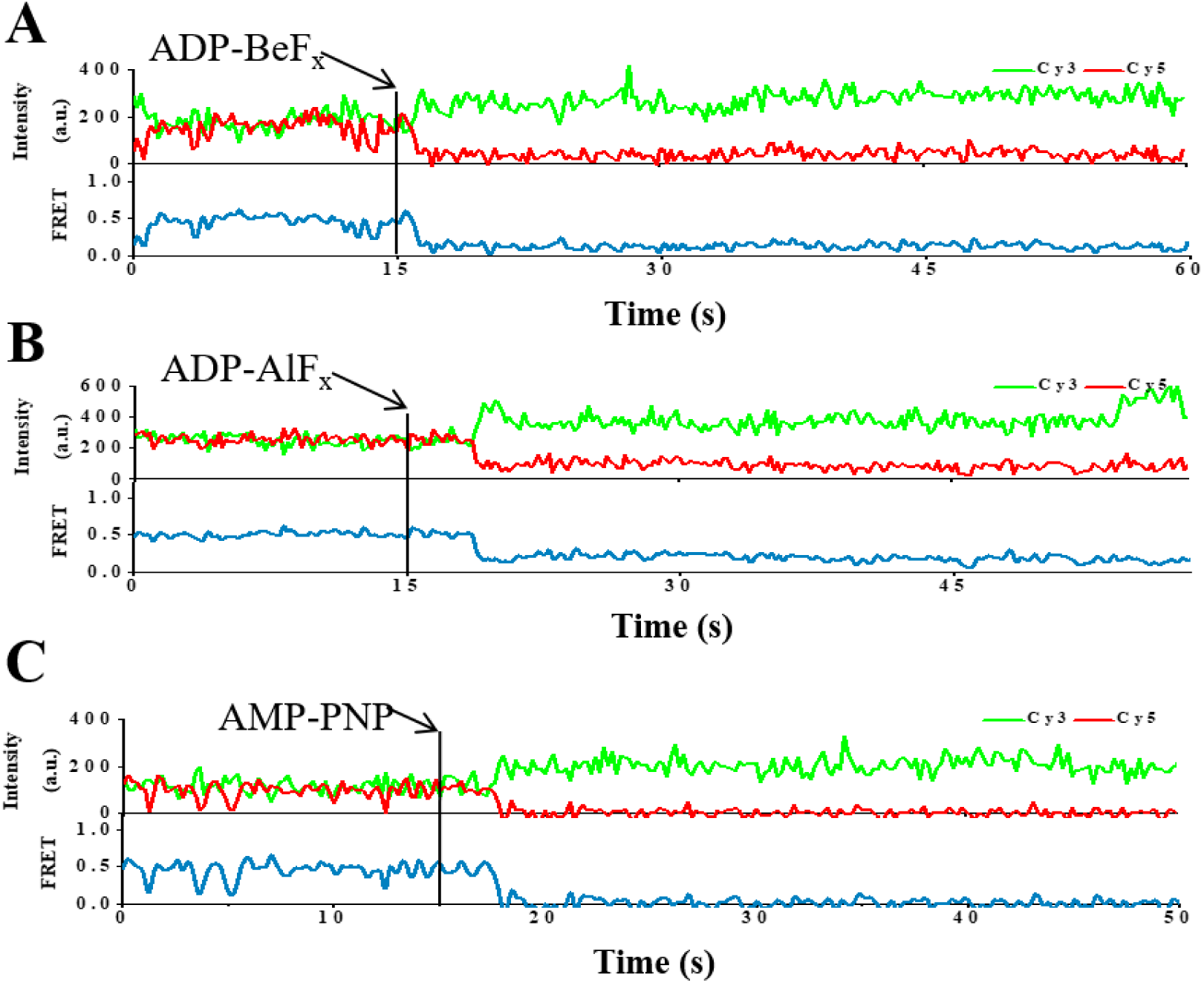
FRET down spikes on the entry side in the presence of various ATP analogs. Stable FRET down spikes were observed in the presence of (A) 1mM ADP-BeF_x_ (1mM ADP, 5mM BeCl_2_, 25mM NaF), (B) 1mM ADP-AlF_x_ (1mM ADP, 5mM AlCl_3_, 25mM NaF), or (C) 1mM AMP-PNP.

**Figure S18.**
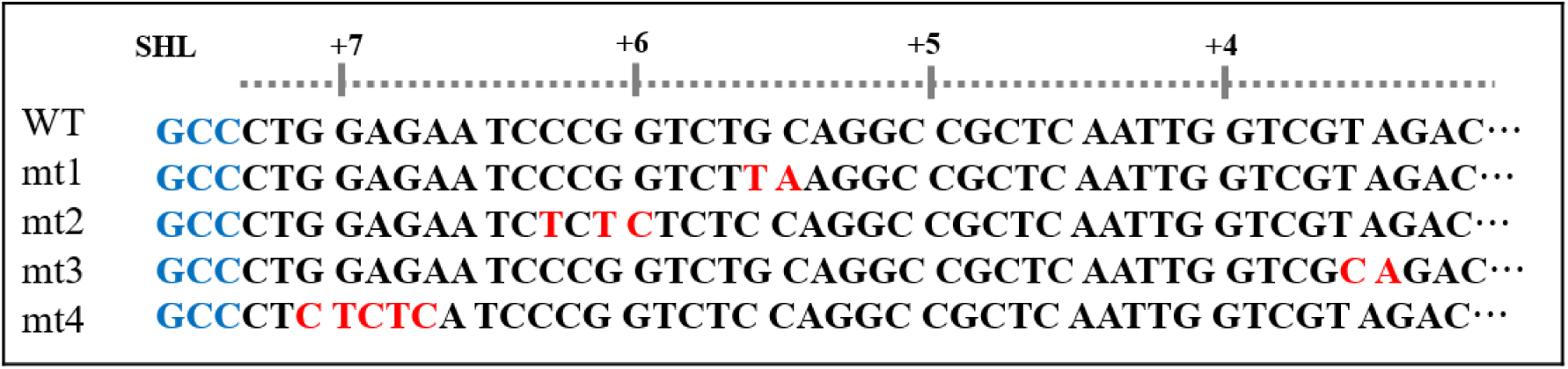
DNA sequences used to study the correlation between DNA unwrapping on the exit side and nucleosome remodeling.

